# Genetic Variants of Phospholipase C-γ2 Confer Altered Microglial Phenotypes and Differential Risk for Alzheimer’s Disease

**DOI:** 10.1101/2022.12.08.519685

**Authors:** Andy P. Tsai, Chuanpeng Dong, Peter Bor-Chian Lin, Adrian L. Oblak, Gonzalo Viana Di Prisco, Nian Wang, Nicole Hajicek, Adam J. Carr, Emma K. Lendy, Oliver Hahn, Micaiah Atkins, Aulden G. Foltz, Jheel Patel, Guixiang Xu, Miguel Moutinho, John Sondek, Qisheng Zhang, Andrew D. Mesecar, Yunlong Liu, Brady K. Atwood, Tony Wyss-Coray, Kwangsik Nho, Stephanie J. Bissel, Bruce T. Lamb, Gary E. Landreth

**Affiliations:** Stark Neurosciences Research Institute, Indiana University School of Medicine, Indianapolis, IN, USA. (A.P.T); (P.B.L); (A.L.O); (G.V.D.P); (J.P); (G.X); (M.M); (B.K.A); (S.J.B); (B.T.L); (G.E.L).; Department of Medical and Molecular Genetics, Center for Computational Biology and Bioinformatics, Indiana University School of Medicine, Indianapolis, IN, USA. (C.D); (Y.L).; Department of Radiology & Imaging Sciences, Indiana University School of Medicine, Indianapolis, IN, USA. (A.L.O); (N.W); (K.N).; Department of Pharmacology and Toxicology, Indiana University School of Medicine, Indianapolis, IN, USA. (G.V.D.P); (B.K.A).; Department of Pharmacology, The University of North Carolina at Chapel Hill, Chapel Hill, NC, USA. (A.J.C); (N.H); (J.S); (Q.Z).; Division of Chemical Biology and Medicinal Chemistry, Eshelman School of Pharmacy, University of North Carolina at Chapel Hill, Chapel Hill, NC, USA. (A.J.C); (Q.Z).; Department of Biochemistry, Purdue University, West Lafayette, IN, USA. (E.K.L); (A.D.M).; Wu Tsai Neurosciences Institute, Stanford University School of Medicine, Stanford, CA, USA. (A.P.T); (O.H); (M.A); (A.G.F); (T.W.C).; Department of Medical and Molecular Genetics, Indiana University School of Medicine, Indianapolis, IN, USA. (S.J.B); (B.T.L).; Department of Anatomy, Cell Biology and Physiology, Indiana University School of Medicine, Indianapolis, IN, USA. (G.E.L).

**Keywords:** Microglia, Alzheimer’s disease, Phospholipase C-gamma-2, P522R variant, M28L variant, Amyloid pathology, Microglia-plaque interactions, Microglial uptake capacity, Synaptic function, transcriptional programs.

## Abstract

Genetic association studies have demonstrated the critical involvement of the microglial immune response in Alzheimer’s disease (AD) pathogenesis. Phospholipase C-gamma-2 (PLCG2) is selectively expressed by microglia and acts in many immune receptor signaling pathways. In AD, PLCG2 is induced uniquely in plaque-associated microglia. A genetic variant of *PLCG2*, PLCG2^P522R^, is a mild hypermorph that attenuates AD risk. We report the identification of a *PLCG2* variant, *PLCG2^M28^*^L^, associated with loss-of-function and confers increased AD risk. PLCG2^P522R^ attenuates disease in an amyloidogenic murine AD model, whereas *PLCG2*^M28L^ exacerbates the plaque burden associated with altered phagocytosis and Aβ clearance. The variants bidirectionally modulate disease pathology by inducing distinct transcriptional programs that identify microglial subpopulations associated with protective or detrimental phenotypes. In summary, these findings identify PLCG2^M28L^ as a new AD risk variant and demonstrate that PLCG2 variants can differentially orchestrate microglial responses in AD pathogenesis that can be therapeutically targeted.

**Graphical abstract:** 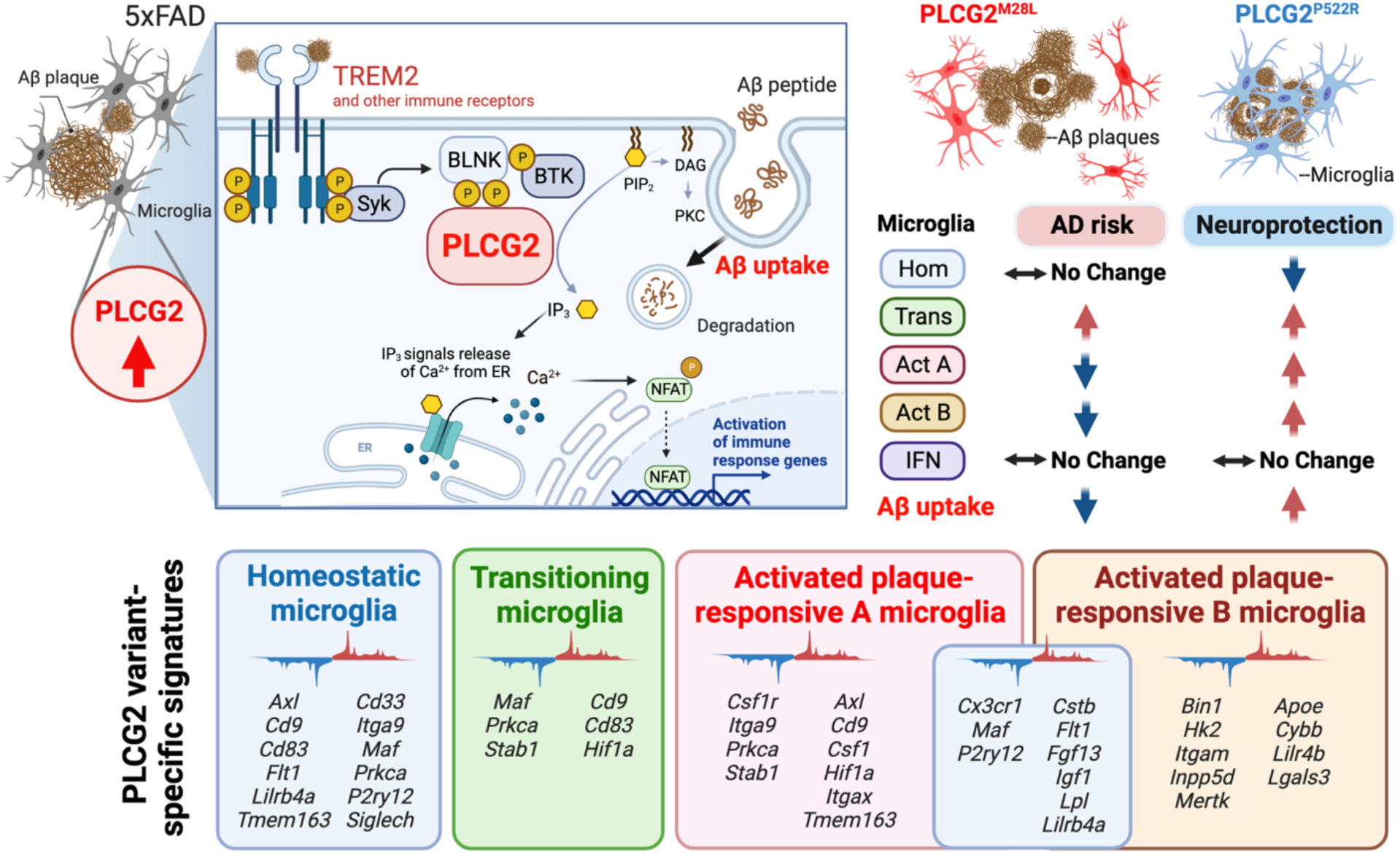

**Highlights:**

- A genetic variant of PLCG2, M28L, is associated with an increased risk for Alzheimer’s disease (AD)
- In an amyloidogenic AD mouse model, PLCG2M28L exacerbates disease pathogenesis
- Conversely, PLCG2P522R, a protective PLCG2 variant, attenuates AD pathogenesis
- The PLCG2 variants uniquely alter the microglial transcriptome and phenotypes

## Introduction

A concerted effort to identify genes that confer altered risk for Alzheimer’s disease (AD) has resulted in the recognition that many disease-risk genes are associated with the microglia-mediated immune response in the AD brain (Keren-Shaul et al., 2017; Masuda et al., 2019; Olah et al., 2020). These studies have provoked a renewed interest in immune mechanisms as therapeutic targets in AD, specifically genes encoding proteins involved in immune receptor signal transduction pathways (Lewcock et al., 2020). Phospholipase C-gamma-2 (PLCG2) is a crucial signaling element employed by various immune receptors that is expressed principally by microglia in the brain. PLCG2 is a key regulatory hub gene for immune signaling (Andreone et al., 2020). Importantly, a PLCG2 gain-of-function variant, PLCG2^P522R^, has been shown to confer reduced AD risk (Sims et al., 2017). However, the role of PLCG2 in AD pathogenesis remains unclear.

PLCG2 is activated by tyrosine phosphorylation, principally by Bruton’s tyrosine kinase (BTK), following ligand binding to cell surface immune receptors, including triggering receptor expressed on myeloid cells 2 (TREM2). Alternatively, PLCG2 can be activated by association with activated forms of Rac GTPases (Bunney et al., 2009; Walliser et al., 2015). The tyrosine phosphorylation of PLCG2 induces a conformational change that displaces its autoinhibitory domain, activating its enzymatic activity. PLCG2 is recruited to the plasma membrane through pleckstrin homology (PH) domains and incorporated into a receptor-associated signaling complex (Falasca et al., 1998). This translocation of PLCG2 to the plasma membrane allows its interaction with its membrane-associated substrate, 1-phosphatidyl-1D-myo-inositol 4,5-bisphosphate (PIP2). PIP2 hydrolysis yields the cytoplasmic secondary messengers, 1D-myo-inositol 1,4,5-trisphosphate (IP3) and diacylglycerol (DAG). The release of IP3 results in the elevation of intracellular calcium levels, promoting the activation of several calcium-regulated transcription factors, including nuclear factor kappa B (NFKB) and nuclear factor of activated T-cells (NFAT) (Schulze-Luehrmann and Ghosh, 2006). In parallel, DAG promotes the activation of numerous downstream signaling cascades, ultimately regulating the cellular immune response (Jing et al., 2021). Importantly, PLCG2 expression is increased in several brain regions in AD patients and the well-studied 5xFAD mouse model of amyloid pathology (Tsai et al., 2022a). A PLCG2 co-expression network analysis using microglial single-cell RNA-seq data identified the association of PLCG2 with inflammatory response-related pathways, consistent with studies documenting its critical involvement in the microglial immune response (Tsai et al., 2022a).

Numerous genome-wide association studies (GWAS) have identified an exonic variant of PLCG2, PLCG2^P522R^, associated with reduced risk for AD (OR=0.68, p=5.38E-10) (Romero-Molina et al., 2022; Sims *et al*., 2017), Lewy body dementia (LBD) and frontotemporal dementia (FTD); this variant is also associated with increased longevity (van der Lee et al., 2019). Among patients with mild cognitive impairment (MCI), PLCG2^P522R^ carriers exhibit a slower cognitive decline rate associated with reduced total tau and phospho-tau levels in the cerebrospinal fluid (CSF) (Kleineidam et al., 2020). Knock-in PLCG2^P522R^ mice show modestly increased basal phospholipase activity (Takalo et al., 2020). Macrophages from these mice exhibit improved survival and viability, increased basal phagocytic activity, and, unexpectedly, elevated cytokine secretion levels (Takalo et al., 2020). PLCG2^P522R^ mice also exhibit an altered microglial gene expression profile, notably involving genes linked to PLC signaling and pathways related to survival, proliferation, and inflammatory responses (Takalo et al., 2020). Paradoxically, reduced expression of genes associated with the phagocytosis of fungal and bacterial particles and enhanced endocytosis of β-amyloid (Aβ) oligomers and dextran is observed in these mice (Maguire et al., 2021). Andreone et al. knocked out *PLCG2* from iPSC-derived microglia and demonstrated that PLCG2 is required for downstream signaling from TREM2, leading to increased microglial viability, phagocytosis and cholesterol metabolism (Andreone et al., 2020). The PLCG2^P522R^ variant has been shown to promote cholesterol metabolism more effectively than the wild-type enzyme, consistent with the view that this variant is hypermorphic (Andreone et al., 2020; Takalo et al., 2020). Furthermore, PLCG2 acts broadly to transduce signals from many immune receptors and is independent of TREM2 (Andreone et al., 2020; Obba et al., 2015; Xu et al., 2009).

Mutant forms of PLCG2 in diverse immune cell populations have been associated with peripheral immune disorders, including leukemias and lymphomas. Specifically, genomic deletions (exon 19 or exons 20 through 22) or somatic mutations (R665W or S707Y) within the regulatory domain of PLCG2 resulted in constitutive activation of PLCG2 and resistance to BTK inhibitor treatment in leukemia patients (Ombrello et al., 2012; Woyach et al., 2014). Surprisingly, through genetic data analysis (Kunkle et al., 2019), we identified the missense PLCG2^M28L^ variant (rs617749044) associated with elevated AD risk, which was previously found in patients with BTK inhibitor-resistant forms of chronic lymphocytic leukemia (Walliser et al., 2016). BTK inhibition in microglia arrests PLCG2 activation and subsequently inhibits phagocytosis (Keaney et al., 2019), suggesting a critical role for PLCG2 in Aβ clearance and AD pathogenesis. Thus, understanding how the PLCG2 variants affect the immune system and their impact on the brain has broad implications in the aging brain, AD and other neurodegenerative disorders.

## Results

### The PLCG2^M28L^ variant is associated with elevated AD risk and reduced PLCG2 expression

Through analysis of data from a large-scale GWAS of >94,000 individuals with late-onset AD originally reported by (Kunkle et al., 2019), we identified a PLCG2 missense mutation (rs617749044) associated with elevated AD risk (OR=1.164; p=0.047) encoding the PLCG2^M28L^ variant. The M28L variant was also identified in a reanalysis of a European-American cohort database and associated with increased risk for AD (OR=1.25 p=0.054) (Olive et al., 2020). In contrast to the M28L variant, the PLCG2 missense exonic variant encoding PLCG2^P522R^ (rs72824905) was protective against AD (OR=0.76, p=0.007) (Kunkle et al., 2019). This finding was validated in a European-American cohort (OR=0.63, p=0.002) (Olive et al., 2020) and was congruent with previous reports (Magno et al., 2021; Sims et al., 2017), verifying that PLCG2^P522R^ is a protective factor in AD.

Structurally, the P522 amino acid residue is positioned between an atypical split PH domain and the c-SH2 domain. The M28 mutation is positioned in the N-terminal PH domain, a domain required for membrane localization (Figures 1A and 1B). We generated mice with the risk-conferring PLCG2^M28L^ variant or the protective PLCG2^P522R^ variant and crossed these mice with 5xFAD mice, an amyloidogenic murine model of AD in which five familial AD-associated mutations are expressed (Oakley et al., 2006), to evaluate the impact of PLCG2 variants on AD pathogenesis. First, we characterized the effect of the variants on PLCG2 expression in the cortex of 5xFAD mice expressing PLCG2^M28L^ (5xFAD^M28L^) or PLCG2^P522R^ (5xFAD^P522R^). The PLCG2 variants did not significantly alter mRNA levels in the brain compared to control mice. However, *Plcg2* gene expression levels were significantly altered in the comparison between 5xFAD^M28L^ and 5xFAD^P522R^ mice (Figure 1C). We found that PLCG2 protein expression levels were significantly reduced in the brains of 5xFAD^M28L^ mice (0.47-fold) compared to those in 5xFAD mice. A significant difference in PLCG2 protein expression levels was observed between 5xFAD^M28L^ and 5xFAD^P522R^ mice (Figure 1D). A reduction in PLCG2^M28L^ protein expression levels was also observed in the spleens of 5xFAD^M28L^ mice (Figure 1E). These data suggest that the PLCG2^M28L^ variant may be less stable and subject to higher turnover than the common variant owing to the conformation change imparted by the mutation and altered enzyme topology or function that subserves the altered AD risk and progression.

**Figure 1.**
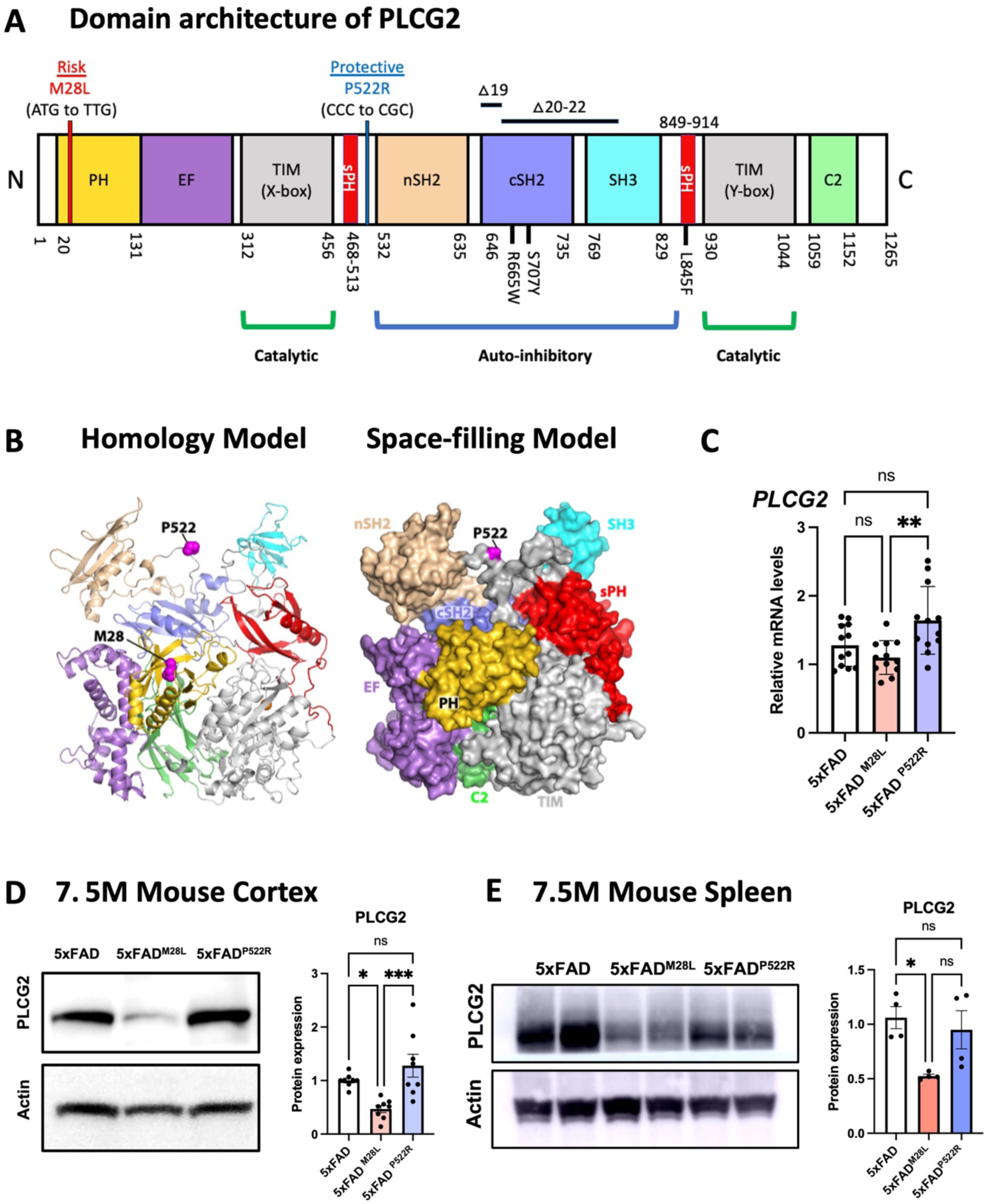
The PLCG2^M28L^ variant is associated with AD risk and downregulates PLCG2 expression. (A) Genetic linkage data of PLCG2^M28L^ and PLCG2^P522R^ with respect to AD risk are shown with the domain architecture of PLCG2 drawn to scale. Somatic mutations (R665W and S707Y) in PLCG2 are shown in the domain architecture. (B) PLCG2^M28L^ (risk) and PLCG2^P522R^ (protective) variants are mapped onto the structure of PLCG2 (magenta spheres) in both the homology model (left) and the space-filling model (right). (C). Gene expression levels of *Plcg2* were assessed in cortical lysates from 7.5-month-old 5xFAD, 5xFAD^M28L^, and 5xFAD^P522R^ mice (n=12 per group, 6 male and 6 female mice). (D) Representative immunoblots and quantifications of PLCG2 protein expression in cortical lysates shows reduced PLCG2 expression in 5xFAD^M28L^ mice (n=8 per group, 4 male and 4 female mice). (E) Representative immunoblots and quantification of PLCG2 protein expression from spleen show reduced PLCG2 expression in 5xFADM28L mice (n=4 per group, 2 male and 2 female mice). All data are presented as the mean ± SEM. * P <0.05; ns: not significant. OR *odds ratio*, N *amino-terminus*, C *carboxyl-terminus*, PH pleckstrin homology domain, EF *EF hand motif*, TIM *TIM barrel*, sPH *split PH domain*, nSH2 *n-terminus, Src Homology 2 domain*, cSH2 *c-terminus Src Homology 2 domain*, SH3 *SRC Homology 3 domain*, C2 *C2 domain*, WT *wild-type*,

### PLCG2 variants affect plaque pathology and microglial uptake of Aβ aggregates

High-resolution T2*-weighted magnetic resonance imaging (MRI) was conducted using a 9.4T/30 MRI scanner to assess amyloid deposition in the brains (Figure 2A). Diffuse plaques were detected by immunofluorescence using an anti-Aβ antibody, 6E10, and compact plaques were visualized by X34 staining (Figure 2B). We observed exacerbated plaque deposition in the brain of 5xFAD^M28L^ compared to that in 5xFAD mice; 5xFAD^M28L^ mice demonstrated a significant elevated hypointense MRI signals in the cortex (1.31-fold) and significantly increased diffuse 6E10-positive and compact X34-stained amyloid deposits in the subiculum. In contrast, 5xFAD^P522R^ mice exhibited a significant reduction in hypointense MRI signals in the cortex (0.77-fold) and 6E10-positive amyloid deposits in the subiculum, demonstrating attenuation of amyloid pathology. Notably, significant differences in 6E10-positive and X34-positive amyloid deposits were observed between 5xFAD^M28L^ and 5xFAD^P522R^ mice (Figure 2C). All genotypes had more diffuse 6E10-positive plaques than compact, strongly X34-positive plaques (Figure 2C). Moreover, compared to 5xFAD^M28L^ mice, 5xFAD^P522R^ mice demonstrated reduced 6E10-positive plaque number and size (Figure S1A) and reduced X34-positive plaque size (Figure S1B). The distribution of Aβ plaques was shifted with a lower overall plaque burden (both diffuse and compact) in the 5xFAD^P522R^ mice compared to that in control and 5xFAD^M28L^ mice (Figure 2D). Moreover, the formation of Aβ plaques was shifted toward more compact (strong X34-positive staining) plaques in the 5xFAD^P522R^ mice and diffuse plaques in the 5xFAD^M28L^ mice compared to that observed in 5xFAD mice (Figure 2E). Collectively, these results establish a functional link between PLCG2 variants that confer elevated or reduced disease risk with amyloid pathology in a murine AD model.

**Figure 2.**
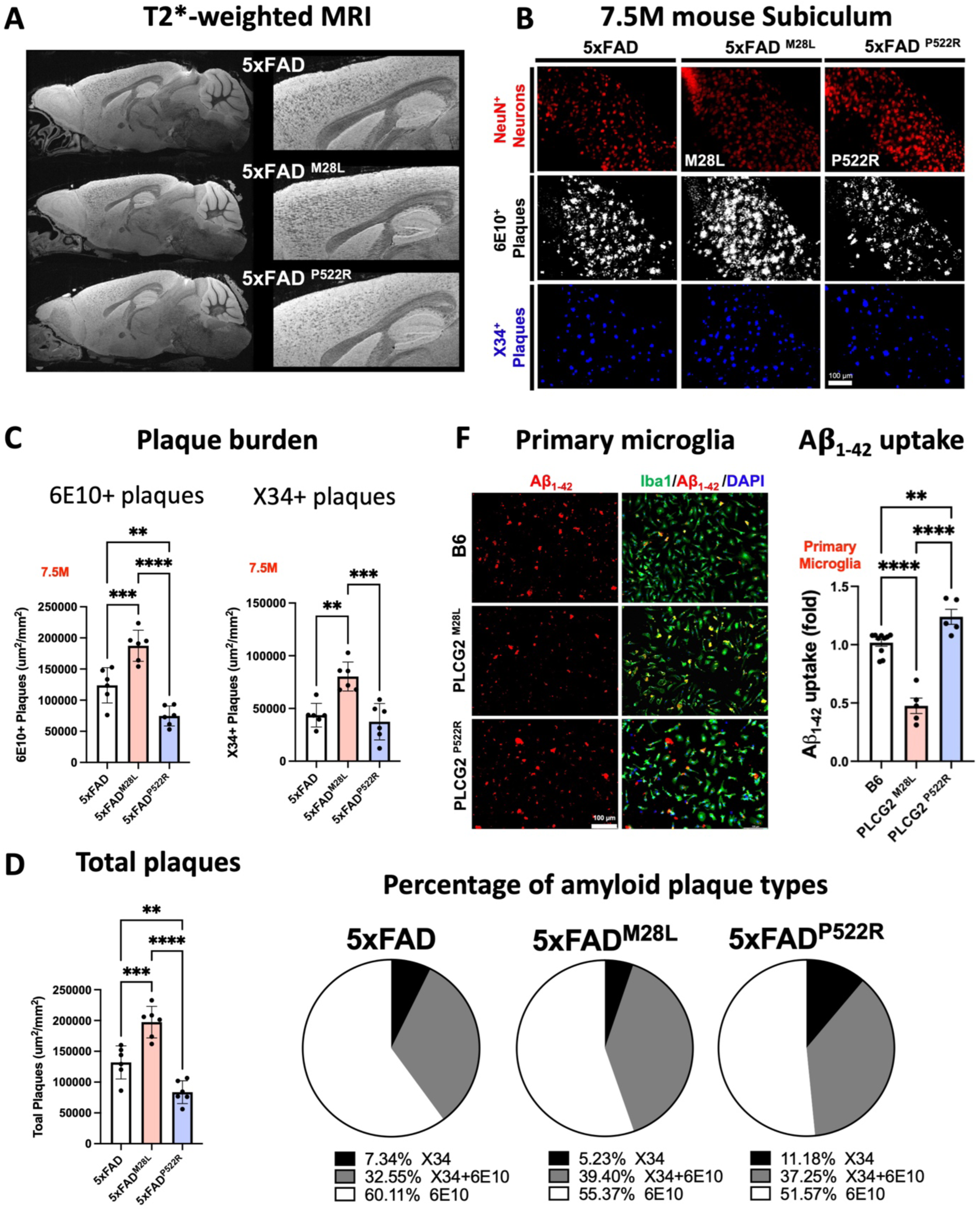
PLCG2 variants affect plaque pathology and microglial uptake of Aβ aggregates. (A) Representative T2*-weighted images of 7.5-month-old AD mice. **(B)** Representative images of amyloid plaques in the subiculum of 7.5-month-old AD mice. **(C)** Immunofluorescence analysis of diffuse 6E10 (white) and compact X34 (blue) positive plaque density. **(D)** Scatter plots show the quantification of the total plaque (6E10-positive and X34-positive) area. **(E)** Graphs denoting the percentage of plaques labeled with X34, 6E10, or their colocalized area. **(F)** Immunofluorescence analysis of primary murine microglia from B6, PLCG2^M28L^, and PLCG2^P522R^ mice incubated with fluorescently labeled-Aβ_1-42_ aggregates (red). Cells were stained with Iba1 (microglia, green) and DAPI (nuclei, blue). Quantification results of Aβ uptake by fluorescence per cell are shown. All data are expressed as the mean values ± SEM (*P < 0.05, **P < 0.01, and ***P < 0.001; ns: not significant).

Microglia play important roles in plaque remodeling and Aβ clearance through phagocytosis (Yuan et al., 2016). Therefore, we examined whether PLCG2 variants affect the intrinsic ability of microglia to take up Aβ. We cultured primary murine microglia from B6, PLCG2^M28L^, and PLCG2^P522R^ mice and incubated the cells with 1.25 μM of HiLyte Fluor 555-labeled Aβ_1-42_ aggregates for 30 min (Figure 2F). The basal uptake of Aβ_1-42_ aggregates was reduced in PLCG2^M28L^ microglia (0.46-fold). Conversely, compared with B6 microglia, the hypermorphic P522R variant increased the microglial uptake of Aβ_1-42_ aggregates (1.22-fold). These results provide supportive evidence that the alteration of AD risk resulting from PLCG2 variants might be due to the microglial function of Aβ_1-42_ uptake in the context of amyloid pathology. These data demonstrate that plaque pathology is particularly sensitive to the PLCG2 variants, exhibiting large and bidirectional changes correlated with their ability to alter disease risk. Importantly, the genotype-dependent alteration in plaque burden is consistent with the effect of the PLCG2 variants on the capacity of microglia to internalize aggregated forms of Aβ that affect plaque remodeling and overall burden.

### PLCG2 variants differentially alter microglial phenotypes and responses to plaques in the 5xFAD mice

The microglia–plaque interaction is a prominent feature of AD pathology; thus, we investigated whether *PLCG2* variants differentially affected microglial responses to plaques in 5xFAD mice. As expected, 7.5-month-old 5xFAD mice exhibited robust microglial clustering around plaques (Figure 3A), reflecting the migration of microglia to deposited Aβ, microglial process envelopment of the plaque and subsequent microgliosis. We did not observe significant changes in the area occupied by ionized calcium-binding adaptor molecule 1 (IBA1; a microglia marker)-positive microglia between genotypes; however, IBA1-positive microglial processes around the plaques were altered by the PLCG2 variants. Within X34-positive plaques, the percent area occupied by PLCG2^M28L^ microglia was significantly reduced compared to that in 5xFAD mice (Figure 3B). Moreover, we observed a similar percent area of C-type lectin domain containing 7A (CLEC7A)-positive microglia, a distinct amoeboid microglia subset (Masuda et al., 2020), in both PLCG2^M28L^ and PLCG2^P522R^ mice. Interestingly, compared to that observed in 5xFAD mice, the percent area of CLEC7A-positive microglia in X34-positive plaques was reduced in PLCG2^M28L^ mice but not in the 5xFAD^P522R^ mice. These findings suggest that plaques induce CLEC7A and PLCG2^M28L^ microglia are less responsive than wild-type (WT) microglia or those expressing PLCG2^P522R^ (Figure 3C). Surprisingly, the numbers of ramified microglia expressing the homeostatic marker purinergic receptor P2Y12 (P2RY12) were significantly increased in 5xFAD^M28L^ mice versus 5xFAD and 5xFAD^P522R^ mice, suggesting an inability or impaired ability of the M28L variant-expressing microglia to transition to a disease-associated microglia phenotype. Although an increase in P2RY12-positive microglia was observed in the 5xFAD^M28L^ mice, we did not find any genotype-dependent differences in P2RY12-positive microglia engaged with X34-positive plaques (Figure 3D).

**Figure 3.**
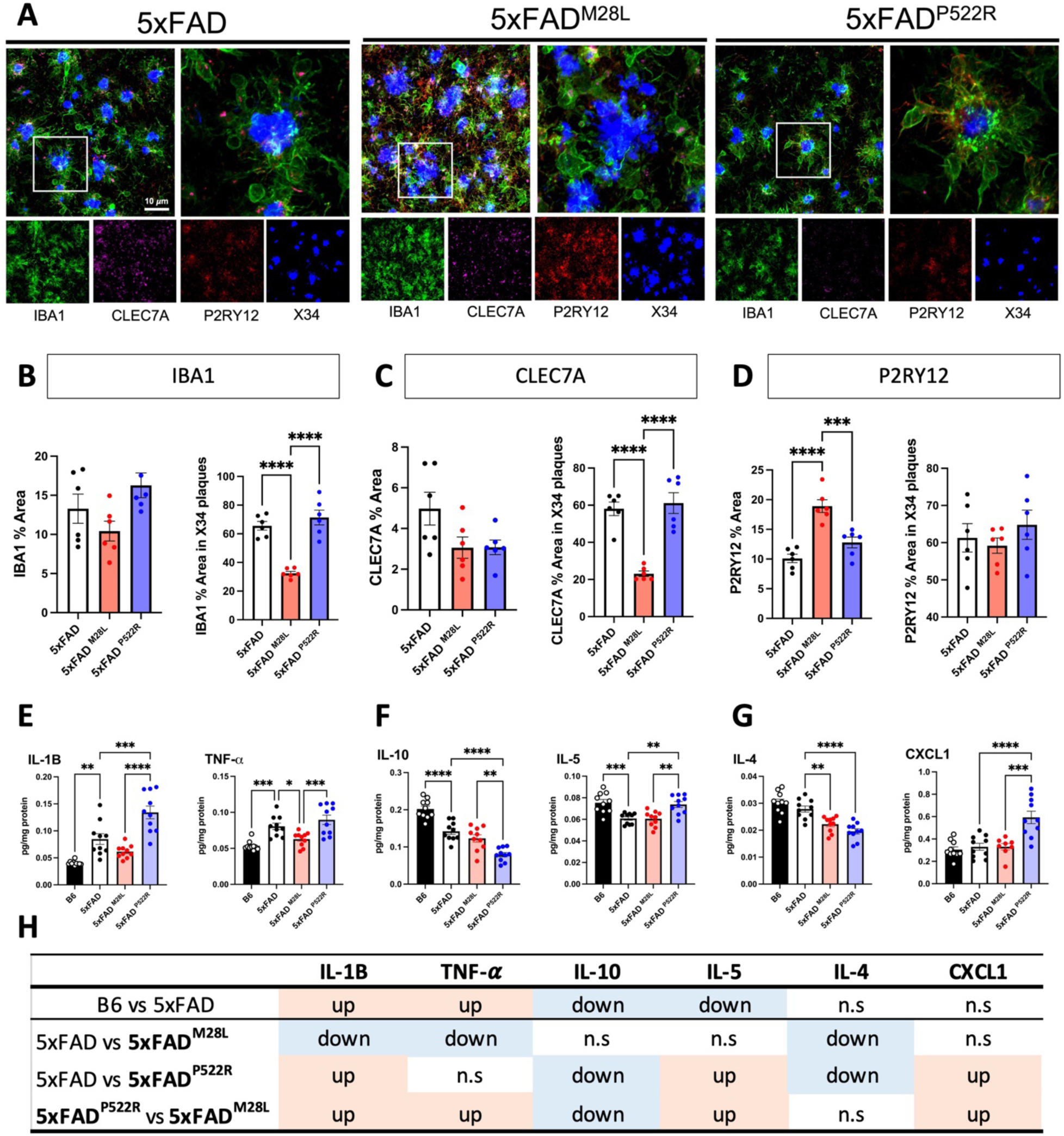
PLCG2 variants differentially alter microglial phenotypes and responses to plaques in the 5xFAD mice. (**A**) Representative images of 7.5-month-old 5xFAD, 5xFAD^M28L^, and 5xFAD^P522R^ mouse subiculum stained with IBA1, CLEC7A, and P2RY12 to label microglia and X34 to label amyloid plaques. Bar, 10 μm. (**B-D**) Scatter plots show quantification of IBA1 (**B**), CLEC7A (**C**), and P2RY12 (**D**) staining in the subicula of 7.5-month-old mice (n=6, 3 male and 3 female mice). The left graphs show the total percentage of the area stained. The right graphs show the quantification of the percentage volume within individual plaque areas. (**E-G**) Protein levels of cytokines were measured from the cortical regions of 7.5-month-old mouse brains (n=10, 5 male and 5 female). (**H**) Table summarizing the results from **E** to **G**. All data were normalized by total protein level. * P <0.05; ** P <0.01; *** P <0.001; **** P <0.0001. IBA1 *Ionized calcium binding adaptor molecule 1*, CLEC7A *C-type lectin domain family 7 member A*, P2RY12 *Purinergic Receptor P2Y12*, TNF-a *Tumor necrosis factor-alpha*, IL *interleukin*, CXCL1 *C-X-C motif chemokine ligand 1*

We next investigated whether the PLCG2 variants resulted in different cytokine levels in the 5xFAD brain using a Meso Scale Discovery (MSD) cytokine panel. Notably, the levels of many cytokines were significantly altered by PLCG2 variants (Figure 3E-G). In the comparison between B6 and 5xFAD mice, the levels of two proinflammatory cytokines, interleukin-1 β (IL-1B), tumor necrosis factor-alpha (TNF-*α*), were increased Figure 2E), and those of two other cytokines, IL-10, IL-5, were decreased (Figure 2F), consistent with previous work documenting that cytokines are influenced by AD pathogenesis (Kinney et al., 2018). Moreover, the levels of IL-1*β*, TNF-*α*, IL-5, and the chemokine (C-X-C motif) ligand 1 (CXCL1) were significantly increased, and levels of IL-10 were reduced in 5xFAD^P522R^ mice versus 5xFAD^M28L^ mice (Figure 3H). These findings suggest that the perturbation of PLCG2-dependent signaling can positively and negatively alter microglial cytokine expression, which contributes to the differential microglial responses to plaques.

### The hypermorphic P522R variant ameliorates impaired synaptic function in 5xFAD mice

It is widely appreciated that cognitive decline correlates with synaptic dysfunction. We next assessed the effect of PLCG2 genotypes on learning and memory in the 5xFAD mouse model, which shows a robust deficit in working memory starting at 4 to 5 months of age (Oakley et al., 2006). At 6 months of age, compared to B6 control mice, both 5xFAD and 5xFAD^M28L^ mice exhibited impaired performance on a Y-maze task; however, we did not observe significant genotype-related differences in working memory impairment between these two strains. Notably, 5xFAD^P522R^ mice were behaviorally similar to B6 control mice in Y-maze task performance, reflecting normal cognition (Figure 4A). These observations suggest that hypermorphic PLCG2^P522R^ plays a protective role against working memory deficits in the amyloidogenic 5xFAD model.

**Figure 4.**
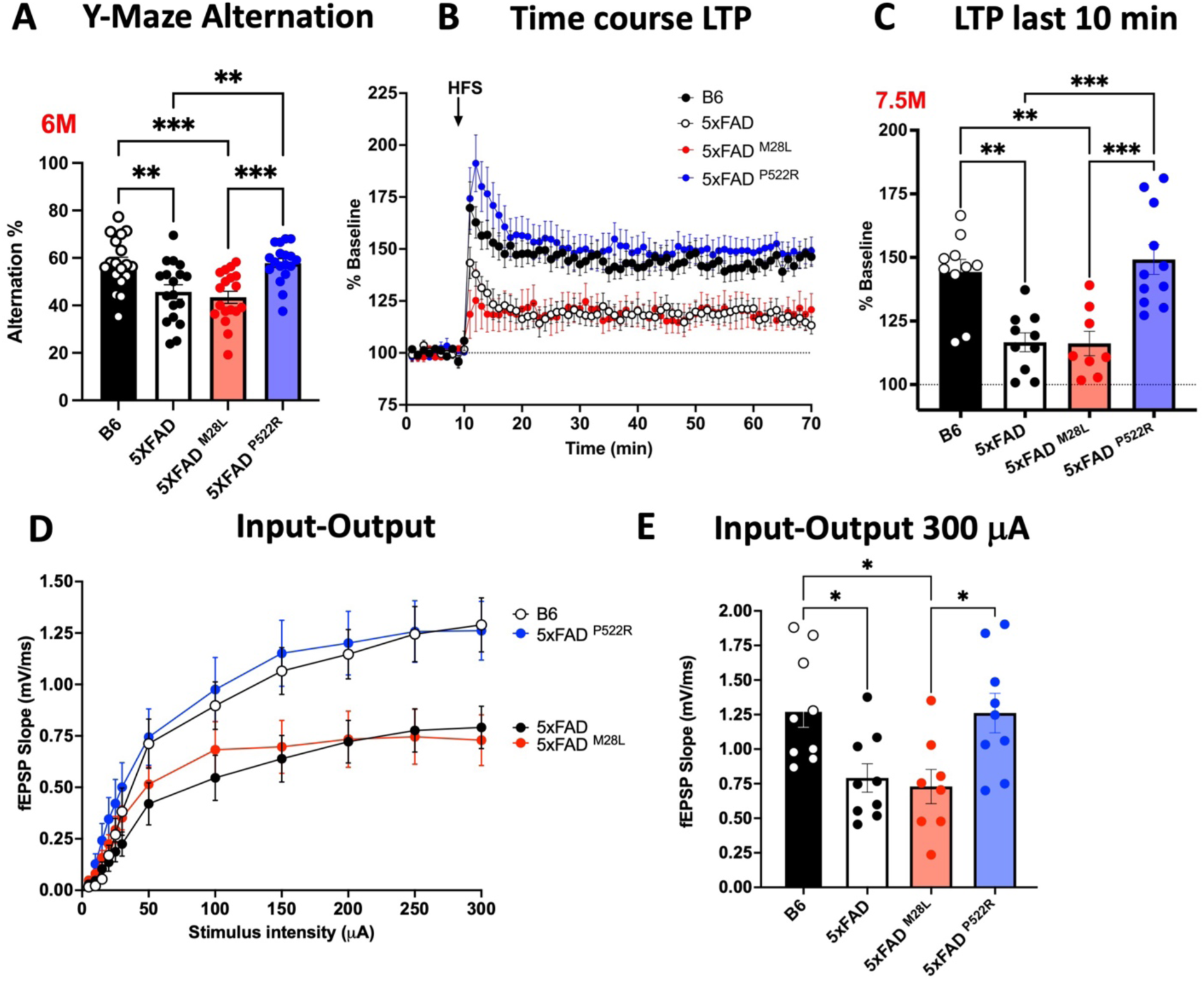
The hypermorphic P522R variant ameliorates impaired synaptic function in 5xFAD mice. **(A)** Working memory of 6-month-old B6, 5xFAD, 5xFAD^M28L^, and 5xFAD^P522R^ mice assessed by percent spontaneous alternation in the Y-maze task (n=18 mice per group, 9 male and 9 female mice). **(B)** The PLCG2^P522R^ variant ameliorated impaired LTP in 7.5-month-old male 5xFAD mice. **(C)** Data show an average of normalized fEPSP slope for the final 10 min of recording (60 to 70 min) relative to 10 min baseline average. **(D)** Input/output curves were obtained by plotting the slope of fEPSPs in the CA1 area of the hippocampus. **(E)** Input/output curves showed diminished basal synaptic transmission in 5xFAD and 5xFAD^M28L^ mice. Statistical analyses were performed by one-way ANOVA followed by Dunnett’s multiple comparison test. The results of individual values from the slices are shown in the scatter plot. Each genotype data set shows at least 4 male mice. All data are expressed as the mean values ± SEM (*P < 0.05, **P < 0.01, and ***P < 0.001). LTP *Long-term potentiation*, fEPSPs *Extracellular recordings of field excitatory postsynaptic potential*, ANOVA *Analysis of variance*

Aβ exposure is associated with impaired synaptic function and blocks long-term potentiation (LTP), a proxy measure of learning and memory (Klyubin et al., 2005). We assessed synaptic plasticity in our mouse models, which showed different severity levels of working memory impairment resulting from the PLCG2 variants. Field recordings of LTP of excitatory postsynaptic potentials (fEPSPs) in hippocampal area CA1 revealed a significantly diminished LTP in the 5xFAD and 5xFAD^M28L^ mice relative to the B6 controls. The LTP was equivalent in the 5xFAD^P522R^ and B6 mice; however, 5xFAD^P522R^ demonstrated greater LTP than 5xFAD and 5xFAD^M28L^ mice, supporting a protective role for PLCG2^P522R^ in preserving synaptic plasticity in 5xFAD mice (Figure 4B and 4C). Next, we measured fEPSP output responses to electrical stimulation. Compared to B6 control mice, output responses were significantly reduced in 5xFAD and 5xFAD^M28L^ mice. 5xFAD^P522R^ mice did not differ from B6 mice and had significantly greater responses than 5xFAD^M28L^ mice (Figure 4D and 4E). We did not find any genotype-related differences in paired-pulse ratio responses, suggesting that presynaptic plasticity was similar among the mouse strains (Figure S2A). Furthermore, we performed whole-cell patch-clamp electrophysiological recordings of hippocampal area CA1 pyramidal neurons to ascertain synaptic differences that could explain the reduced LTP and output responses in 5xFAD and 5xFAD^M28L^ mice. Analysis of spontaneous excitatory postsynaptic currents (sEPSCs) revealed significant effects on sEPSC frequency and sEPSC amplitudes in 5xFAD^P522R^ mice versus 5xFAD or 5xFAD^M28L^ mice. Interestingly, there were no differences in these measures between 5xFAD^P522R^ mice and B6 controls (Figure S2B). We further evaluated differences in excitatory transmission by measuring EPSCs mediated by AMPA and NMDA glutamate receptors. There were significantly smaller AMPA/NMDA receptor current ratios in the 5xFAD and 5xFAD^M28L^ mice relative to the B6 mice, but no difference between the 5xFAD^P522R^ mice and the B6 mice (Figure S2C). In measures of inhibitory transmission, the spontaneous inhibitory postsynaptic current (sIPSC) frequency and amplitudes were significantly lower in 5xFAD and 5xFAD^M28L^ mice relative to the B6 mice. Similarly, the 5xFAD^P522R^ showed a similar pattern to the B6 mice (Figure S2D). Overall, the reduced LTP and input-output response in 5xFAD and 5xFAD^M28L^ mice is likely a result of predominating reduced postsynaptic glutamate receptor responses rather than an expected increased GABA transmission. These findings document a functional role of PLCG2 in governing synaptic functionality, as evidenced by the effect of its hypermorphic variant PLCG2^P522R^. However, we did not detect differences between 5xFAD mice and 5xFAD^M28L^ mice.

### PLCG2 variants elicit distinct transcriptional programs in the 5xFAD mouse brain

PLCG2 is a critical signaling intermediate subserving the actions of several immune receptors that selectively alter the microglial transcriptome (Jackson et al., 2021). We performed gene expression analysis using bulk RNA-seq data from 7.5-month-old AD mice to evaluate the impact of PLCG2 variants on gene expression in the brain. In the comparison between 5xFAD^M28L^ and 5xFAD mice, we identified 47 significant differentially expressed genes (DEGs); 20 upregulated (15 genes with fold changes greater than 1.5) and 27 downregulated (25 genes with fold changes greater than 1.5) (Figure 5A). The pathway analysis identified several Gene Ontology (GO) terms, including “antigen processing and presentation, tumor necrosis factor production, interleukin-1 beta production, inflammatory response, endocytosis and phagocytosis” (Figure 5B), suggesting an important role for PLCG2^M28L^ in inflammation-and endocytosis/phagocytosis-related pathways in AD. In the comparison between 5xFAD^P522R^ and 5xFAD mice, we identified 763 significant DEGs; 568 upregulated (384 genes with fold changes greater than 1.5) and 195 downregulated (61 genes with fold changes greater than 1.5) (Figure 5C). The pathway analysis identified several GO terms, including “multicellular organism metabolic process, glial cell development, regulation of phagocytosis/endocytosis, inflammatory response, and lipid metabolic process” (Figure 5D), supporting our results indicating that *PLCG2* variants play an important role in phagocytosis/endocytosis and inflammatory responses in AD.

**Figure 5.**
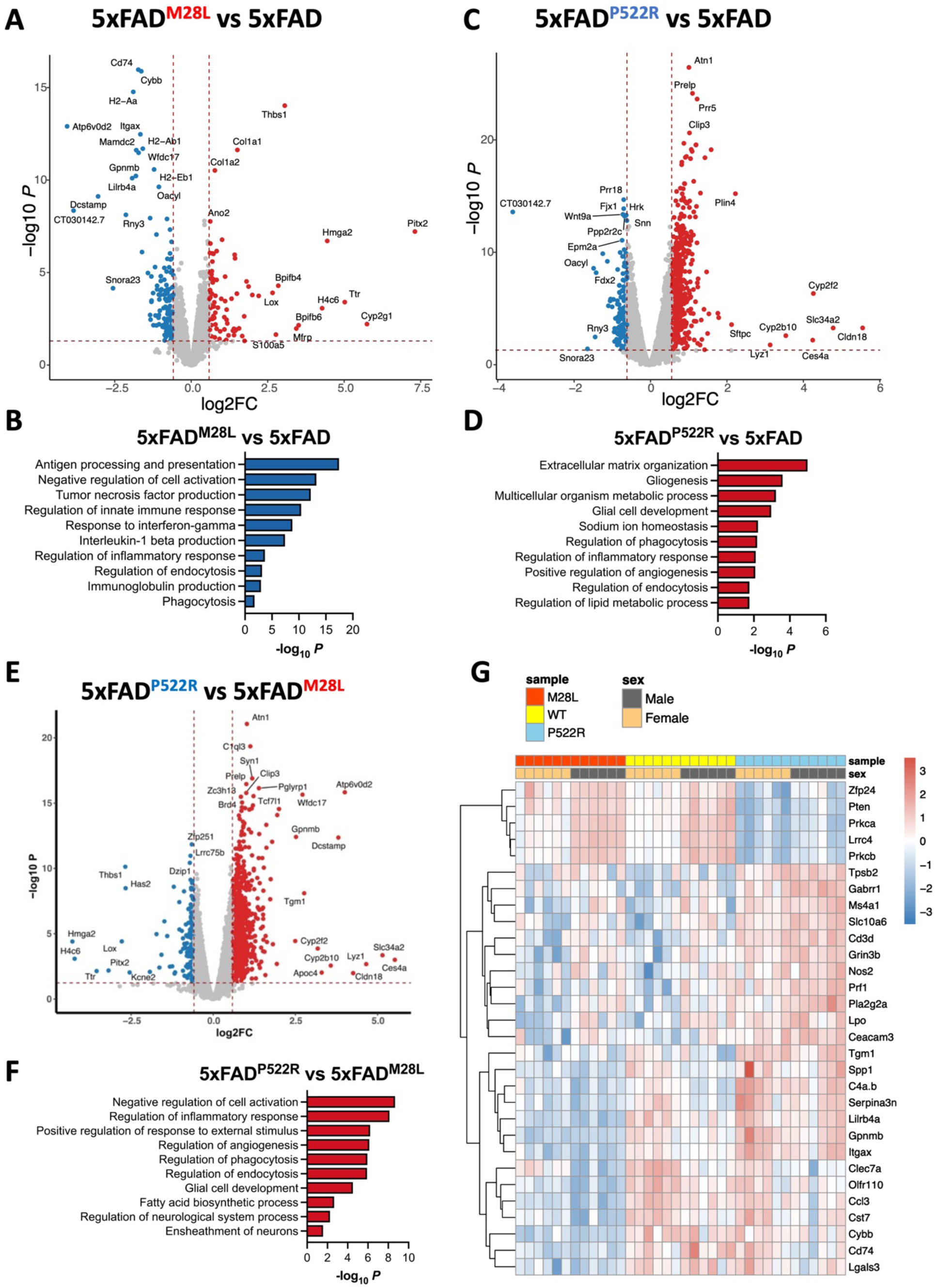
PLCG2 variants elicit distinct transcriptional programs in 5xFAD. Bulk RNA sequencing of 7.5-month-old mice of the indicated genotype was carried out. (**A**) The volcano plot shows significant DEGs (FDR<0.05, FC>1.5) in the cortices from 5xFAD^M28L^ mice (n=8, 4 male and 4 female mice) versus 5xFAD mice (n=8, 4 male and 4 female mice). (**B**) Top 10 Gene Ontology biological processes identified through analysis of the DEGs between 5xFAD^M28L^ and 5xFAD mice. (**C**) The volcano plot shows significant DEGs (FDR<0.05, FC>1.5) in the cortices from 5xFAD^P522R^ mice (n=8, 4 male and 4 female mice) versus 5xFAD mice. (**D**) Top 10 Gene Ontology biological processes identified through analysis of the DEGs between 5xFAD^P522R^ and 5xFAD mice. (**E**) The volcano plot shows significant DEGs in the cortices from 5xFAD^M28L^ mice versus 5xFAD^P522R^ mice. (**F**) Top 10 Gene Ontology biological processes identified through analysis of the DEGs between 5xFAD^M28L^ and 5xFAD^P522R^ mice. (**G**) Nanostring nCounter Gial Profiling and Neuropathology panels were employed to analyze cortical RNA. The gene expression heatmap shows selected DEGs derived from the NanoString analysis of cortices of 7.5-month-old mice (each genotype n=12, 6 male and 6 female). DEGs *differentially expressed genes*, FDR *false discovery rate*, FC *fold change*

We performed differential gene expression analyses using bulk RNA-seq data from 5xFAD^M28L^ and 5xFAD^P522R^ mice to further characterize the differences between the transcriptional programs of the risk and protective variants. We identified 593 DEGs, 439 of which were upregulated (356 genes with fold changes greater than 1.5) and 154 downregulated (61 genes with fold changes greater than 1.5) (Figure 5E). The pathway analysis identified several GO terms, including “regulation of inflammatory response, phagocytosis/endocytosis, glial cell development, fatty acid biosynthesis process and neurological system process” (Figure 5F). These findings indicate that distinct transcriptional programs in 5xFAD mouse brains are elicited by PLCG2 variants, altering amyloid pathogenesis.

Our results demonstrated that PLCG2 variants modulate disease pathology by inducing specific microglial phenotypes and molecular signatures. RNA from the cortices of 7.5-month-old AD mice was analyzed to validate these findings using the nCounter Glial Profiling Panel and Neuropathology Panel from array-based amplification-free NanoString technologies, which comprises 1,266 genes mainly involved in glial biology, neuro-glial interactions, and neurodegeneration. Thirty-four DEGs (all downregulated, 11 with fold changes greater than 1.5) and several genes associated with disease-associated microglia (DAM), including *Itgax*, *Gpnmb*, *Ccl3*, *Cd74*, *Cybb*, *Lgals3*, *Spp1*, and *Lpl* (highlighted in Figure 5G), were found in 5xFAD^M28L^ compared to 5xFAD mice, supporting our findings that PLCG2^M28L^ impaired the ability of microglia to transition into a disease-associated phenotype. Furthermore, 579 DEGs, 350 upregulated (108 genes with fold changes greater than 1.5) and 229 downregulated (71 genes with fold changes greater than 1.5) were found in 5xFAD^P522R^ mice compared to 5xFAD mice. Most of the identified DEGs were related to neuronal connectivity, transmitter response, and structure of axon and dendrite, including *Gabrr1*, *Prl*, *Grin3b*, *Prkca*, *Pten*, *Lrrc4*, and *Cacnb4*, supporting our findings that PLCG2^P522R^ plays a protective role in synaptic function in 5xFAD mice.

### Single nuclei RNA-seq distinguished the cell type–specific effects of PLCG2 variants in the AD brain

We performed snRNA-seq on 7.5-month-old 5xFAD, 5xFAD^M28L^, and 5xFAD^P522R^ mice (4 mice per genotype) to investigate the impact of PLCG2 variants and amyloid pathology across cell types in these AD mice. NeuN-negative nuclei were sorted from the mouse brains to characterize the enrichment of glial cell types and states involved in AD. A total of 62,588 individual nuclei were obtained. Visualization in uniform manifold approximation and projection (UMAP) space separated nuclei into distinct clusters across all samples, which we mapped to 6 major cell types (Figure 6A). These clusters were manually identified based on the expression of known cell-type-specific markers, including oligodendrocytes (Oligo; Cluster 0), oligodendrocyte progenitor cells (OPCs; Cluster 1), microglia (Cluster 2), astrocytes (Ast; Cluster 3), neurons (Neuron; Cluster 4), and endothelial cells (Endo; Cluster 5) (Figure 6B). The distributions of cell types differed among genotypes. The percentages of each cell type are shown in Figure 6C, and the PLCG2 variant-associated microglial signatures were comprehensively analyzed (Figure 7). Notably, only microglial signatures in the brain were differentially altered by the PLCG2 variants, suggesting PLCG2 variants modulate disease pathologies through microglia-specific transcriptional programs.

**Figure 6.**
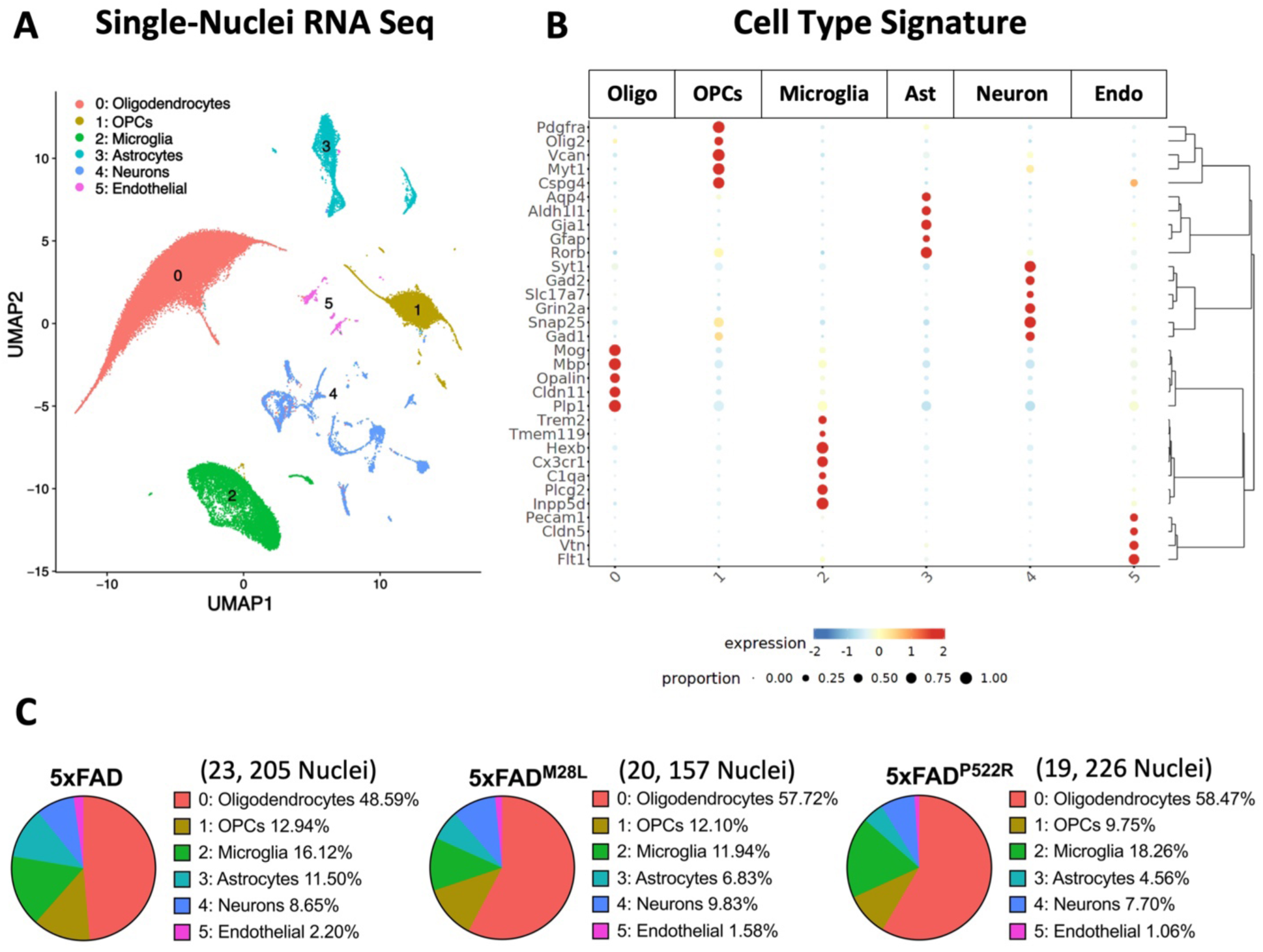
Single nuclei RNA-seq distinguishes the cell type–specific effects of PLCG2 variants -in the AD brain. **(A)** Uniform Manifold Approximation and Projection (UMAP) of 62,588 nuclei captured from 12 cortex samples across three genotypes of AD mice, annotated and colored by cell type. **(B)** Heatmap showing the expression of specific markers in each sample, identifying each cluster in **(C)** Pie chart showing the percentage of clusters in each genotype. Oligo *Oligodendrocytes*, OPCs *Oligodendrocyte progenitor cells*, Ast *Astrocytes*, Endo *Endothelial cells*

**Figure 7.**
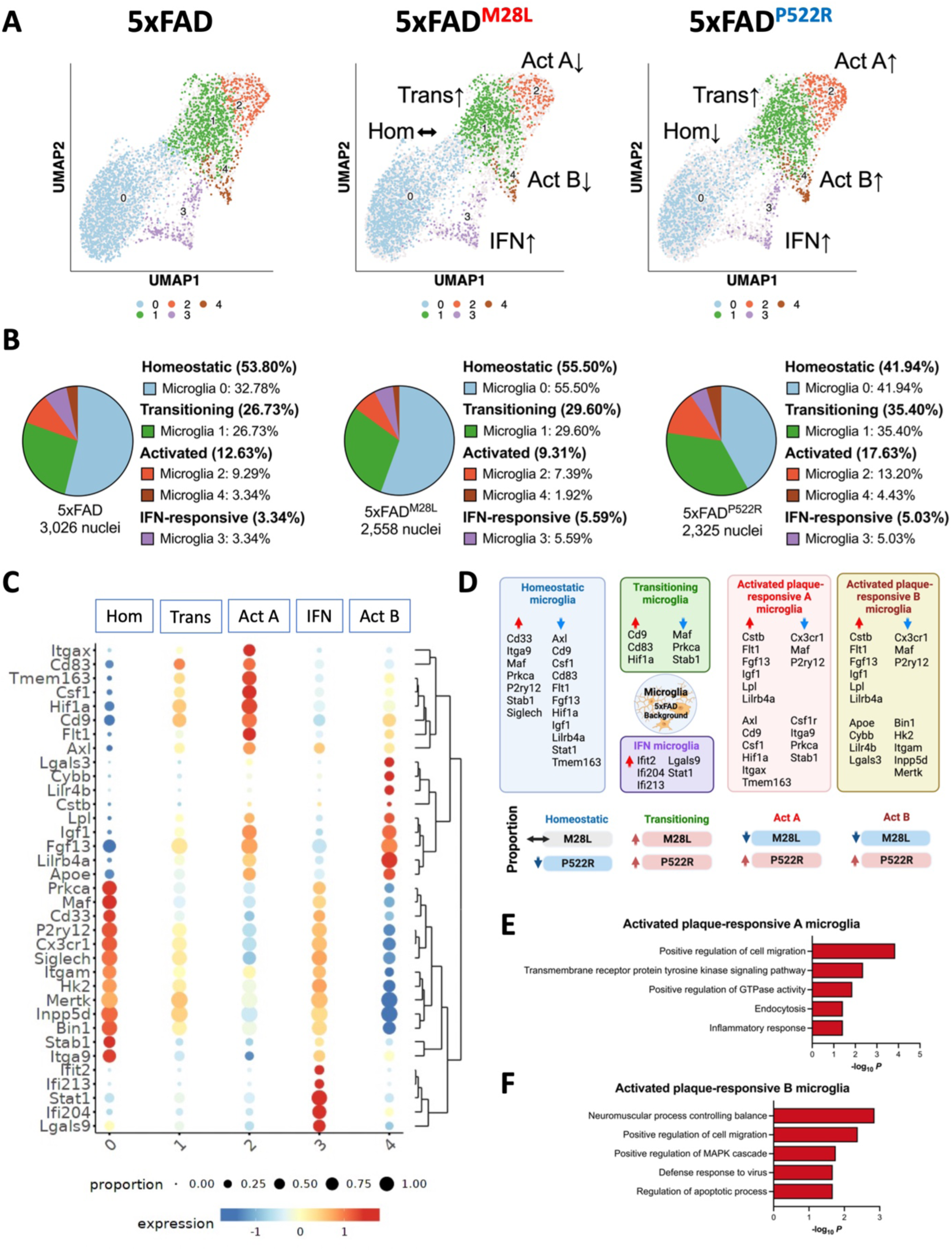
Single nuclei RNA-seq identifies PLCG2 variant-specific microglial signatures in AD **(A)** UMAP plot of 7,909 nuclei showing the re-clustered microglia (from cluster 2 in Figure 6A) annotated and colored by microglial subcluster. **(B)** Pie chart showing the percentage of microglial subclusters. **(C)** Heatmap showing the expression of canonical microglial genes in each microglial subclusters, including homeostatic (Hom), transitioning (Trans), activated plaque-responsive (Act A and Act B), and IFN-responsive (IFN) for each genotype. **(D)** Schematic illustration showing microglial signature altered by PLCG2 variants switching between homeostatic, transitioning, activated plaque-responsive, and IFN-responsive microglial subclusters. Key genes involved in each microglial population are shown. The arrows indicate upregulated (red) or downregulated (blue) genes or proportions. **(E)** Top 5 Gene Ontology biological processes identified through analysis of the upregulated DEGs in activated plaque-responsive microglia A subcluster. **(F)** Top 5 Gene Ontology biological processes identified through analysis of the upregulated DEGs in activated plaque-responsive microglia B subcluster. Hom *homeostatic*, Trans *transitioning*, Act *activated plaque-responsive,* IFN *interferon-responsive*

### Single nuclei RNA-seq identified PLCG2 variant-associated microglial signatures in AD

Robust induction of microglial PLCG2 expression was observed in the brains of 5xFAD mice and patients with AD (Tsai et al., 2022a). We next focused on the 7,909 nuclei mapped in the microglial cluster and examined genes showing differential expression in distinct microglial subpopulations. The analysis revealed 5 discrete microglial clusters, each representing a unique microglial state, including previously characterized homeostatic, transitioning, activated plaque-responsive (Act), and interferon (IFN)-responsive microglial subsets (Keren-Shaul et al., 2017; Krasemann et al., 2017; Olah et al., 2020) (Figure 7A and 7B). Our results demonstrated a subpopulation of microglia, cluster 0, expressing homeostatic microglial genes, such as *P2ry12* and *Cx3cr1* (Figure 7C). In contrast, cluster 2 microglia (termed Act A microglia) expressed more genes upregulated in endocytosis and inflammatory response microglia, including *Itgax*, *Cd9*, and *Axl* (Figure 7C and 7E). Cluster 1 microglia were transitioning microglia, which exhibited slightly increased expression of genes enriched in cluster 2 and slightly decreased expression of genes enriched in cluster 1. Cluster 4 microglia (termed Act B microglia) were enriched in genes related to apoptotic processes, lipid metabolism and plaque compaction, such as *Lpl*, *Apoe*, *Lgals3* and *Lilr4b* (Figure 7C and 7F). In addition, IFN-responsive microglia (cluster 3) demonstrated enrichment of genes such as *Ifit2*, *Ifi204*, and *Ifi213* and were mapped to cluster 3 (Figure 7C). Compared to 5xFAD mice, 5xFAD^M28L^ mice show reduced nuclei percentages in clusters 2 and 4; 5xFAD^P522R^ mice show increased nuclei percentages in clusters 1, 2, and 4 and reduced nuclei percentages in clusters 0 (Figure 7B). 5xFAD mice with the protective PLCG2^P522R^ variant exhibited an increased number of microglia associated with pathways related to cell migration, endocytosis, inflammatory responses, and apoptotic process; these microglia were reduced in 5xFAD mice with the risk PLCG2^M28L^ variant. Our single-nuclei analysis revealed that risk and protective PLCG2 variants elicit differential programming of transcriptomes (Figure 7D) linked to their association with plaques and Aβ clearance leading to opposite effects on disease pathogenesis in 5xFAD mice.

## Discussion

Genetic studies have linked the PLCG2^P522R^ variant with a reduced risk for AD; however, the functionality of PLCG2 in AD pathophysiology remains poorly defined. Notably, we identified the PLCG2^M28L^ variant associated with loss-of-function and elevated AD risk, which allowed us to investigate the mechanisms of PLCG2 variants and identify molecular signatures and pathways discriminating the divergent effects of genetic variants of PLCG2 in AD mouse models. Characterizing specific microglial phenotypes imparted by the PLCG2 AD risk variants allowed the discrimination of microglial mechanisms subserving altered disease risk. Moreover, the phenotypic effects of the variants validate PLCG2 as a critical hub gene that regulates a diverse range of effectors.

The missense mutations appear to have rather modest effects on PLCG2 enzymatic activity. PLCG2^P522R^ exhibits modestly elevated activity (Jing et al., 2021; Magno et al., 2019; Takalo et al., 2020), whereas the activity of the M28L variant was not different from that of the WT enzyme (Walliser et al., 2016). The question of how the variants affect PLCG2 functionality needs to be revisited in a more rigorous analysis of its interactions with other signaling elements.

We found that the expression levels of the PLCG2 variants in the brain differed, likely contributing to the functional differences in the phenotypes observed. The level of expression of PLCG2^P522R^ in the 5xFAD brain was not different from that of the WT enzyme as measured by either mRNA or protein levels (Figures 1C and 1D). However, 5xFAD mice bearing the PLCG2^M28L^ variant exhibited significantly lower protein levels without a measurable change in mRNA levels (Figure 1D). We verified the reduction in protein expression of spleens from these mice (Figure 1E). The reduced levels of PLCG2^M28L^ confer a loss-of-function effect, and the observed phenotypes are consistent with those observed in PLCG2-deficient microglia (Andreone et al., 2020; Jing et al., 2021). The basis of the reduced protein levels of PLCG2^M28L^ is unknown.

PLCG2 is a critical participant in the immune response induced by amyloid plaques in AD patients and animal models (Tsai et al., 2022b). Indeed, PLCG2 is robustly induced in a subset of microglia that are physically associated with plaques. The PLCG2 variants differ in their response to amyloid plaques (Figure 3) with respect to the density of microglia associated with plaques and their ability to remodel plaque size and compaction (Figure 2). The PLCG2^M28L^ variant is associated with an overall greater plaque burden, whereas PLCG2^P522R^ mice have a reduced plaque burden (Figure 2D). Significantly, PLCG2^M28L^ mice demonstrate a substantial reduction in the number of plaque-associated microglia, suggesting that PLCG2^M28L^ microglia may be unable to shift their phenotypes to a more plaque-responsive microglial state, similar to TREM2 loss-of-function microglia, as they cannot efficiently mobilize a robust response to the deposited amyloid (Cheng-Hathaway et al., 2018).

An important feature of the PLCG2^P522R^ variant is its robust effects in sustaining synaptic function and working memory that are impaired in 5xFAD and 5xFAD^M28L^ mice, which provides important information on determinants of brain resilience. Specifically, LTP and glutamatergic and GABAergic transmission are impaired in 5xFAD and 5xFAD^M28L^ mice (Figure 4 and Figure S2). We hypothesized that the loss of LTP is more likely a product of reduced glutamate transmission than enhanced GABA transmission. This hypothesis is consistent with previous studies that revealed that Aβ deposits contribute to disturbed glutamatergic neurotransmission (Chang et al., 2006; Danysz and Parsons, 2012). In our transcriptional analyses, we identified many GO terms associated with synaptic function and structure that support our physiological measures and may provide clues to the mechanistic underpinnings of the altered and protected synaptic function associated with the different variants. These findings also require more directed analyses to determine causative mechanisms. Synaptic function could be protected by the stimulation of a protective microglial response in the PLCG2^P522R^ variant or through PLCG2^P522R^ variant-enhanced phagocytic microglial actions associated with plaque remodeling and synapse pruning. Overall, our findings indicate that the related role of the PLCG2^P522R^ variant in mediating excitatory and inhibitory synaptic function might be important for determining the mechanisms by which microglia modulate disease progression. However, the molecular signatures of neuronal populations among genotypes in AD mice and the exact mechanism by which PLCG2 variants regulate synaptic function in the absence of amyloid plaques require further assessment.

PLCG2 regulates the expression of genes governing pathways related to inflammation, lipid metabolism, microglial viability and phagocytosis (Takalo et al., 2020; Tsai et al., 2022a). However, the effects of its inactivation or genetic variants on transcriptional programming in AD have yet to be extensively explored in AD animal models. It is of particular interest to determine the unique microglial subpopulations that are differentially affected by PLCG2 genotypes and linked to altered disease risk. Our single-nuclei RNA-seq analysis data indicated that PLCG2 variants regulate the composition of microglial clusters in the AD brain. These results were consistent with our previous analysis of AD chimeric mice with xenografted human iPSC-derived microglia expressing PLCG2^P522R^ (Claes et al., 2022), in which the proportions of microglia enriched in *Spp1*, *Fabp5*, and *Cd74* were elevated. Moreover, 5xFAD^P522R^ mice exhibit a plaque-responsive microglial subcluster enriched in *Hcar2*, a microglial receptor we recently identified to be required for efficient and neuroprotective microglial responses to amyloid pathology (Moutinho et al., 2022).

In the current study, we identified specific microglial subclusters enriched in phagocytic and plaque-responsive genes conferring both reduced and elevated AD risk that enabled us to identify distinct stages in the plaque-responsive microglial phenotypes associated with the PLCG2 variants. Initially, plaque deposition and microglia recruitment results in the activation of a set of genes, including *Cd9*, *Cd83*, and *Hif1a*, within a subset of microglia; concomitantly, a set of homeostatic genes is downregulated in non-plaque-associated microglia, such as *Maf*, *Prkca* and *Stab1*, shifting the microglial phenotype from cluster 0 (homeostatic) to cluster 1 (transitioning). The second phase of plaque-responsive microglia activation includes the induction of lipid metabolism or phagocytic disease-associated pathways (e.g., *Cstb*, *Flt1*, *Fgf13*, *Igf1*, *Lpl*, and *Lilrb4a*), shifting the microglial phenotype from cluster 1 (transitioning) to cluster 2 (activated plaque-responsive A; Act A) or cluster 4 (activated plaque-responsive B; Act B). Compared to 5xFAD mice, 5xFAD^M28L^ mice with increased amyloid burdens have increased percentages of transitioning microglia; however, the phenotypes were not shifted to a more activated stage compared to that of activated plaque-responsive microglia (Act microglia). In contrast, 5xFAD^P522R^ mice exhibit reduced percentages of microglia-enriched homeostatic markers and increased percentages of transitioning and Act microglia. Notably, our studies identified two types of Act microglia (Act A and Act B; Figures 6C and 6D). Nuclei mapped to Act A microglia were enriched in genes related to endocytosis and inflammatory response and highly expressed *Axl*, *Cd9*, *Csf1*, *Hif1a*, *Itgax*, and *Tmem163*. Moreover, nuclei mapped into Act B microglia were enriched in genes related to migration and apoptotic processes and highly expressed *Apoe*, *Cybb*, *Lilr4b*, and *Lgals3.* Together, our results shed new light on the role of PLCG2 variants in microglial processes associated with AD risk and amyloid pathologies.

In summary, our results validate the human genetic findings that loss-of-function and gain-of-function variants of the same gene, *PLCG2*, have opposite effects on transcriptional mechanisms, conferring altered microglial phenotypes and differential pathogenesis in AD mice. The data are consistent with the conclusion that PLCG2^M28L^ is a loss-of-function variant due to the reduced expression of PLCG2, leading to an impaired microglial response to plaques, suppressed cytokine release, downregulated plaque-associated and disease-associated microglial genes, and increased plaque deposition. Conversely, PLCG2^P522R^ appears to be a mild hypermorph that leads to a broad range of positive effects in 5xFAD mice, including a reduction in plaque burden, amelioration of impaired synaptic function, and rescue of working memory deficits, indicating that promoting a neuroprotective microglial response to amyloid pathology could limit AD progression. Overall, our studies suggest that PLCG2-directed therapeutics may provide novel strategies to induce neuroprotective microglial responses to attenuate AD pathogenesis.

## STAR methods

### Whole-genome sequencing (WGS)

Pre-processed whole-genome sequencing data were obtained from the Accelerating Medicines Partnership for Alzheimer’s Disease (AMP-AD) Consortium (https://adknowledgeportal.synapse.org/Explore/Programs/DetailsPage?Program=AMP-AD) through the Synapse database (https://www.synapse.org/) (Tsai et al., 2022a). Whole-genome sequencing libraries were prepared using the KAPA Hyper Library Preparation Kit per the manufacturer’s instructions. Libraries were sequenced on an Illumina HiSeq X sequencer using paired-end read chemistry and read lengths of 150bp. The paired-end 150bp reads were aligned to the NCBI reference human genome (GRCh37) using the Burrows-Wheeler Aligner (BWA-MEM) (Li and Durbin, 2010) and processed using the GATK best practices workflow that marks potential duplicates, locally realigns any suspicious reads, and re-calibrates the base-calling quality scores using Genome Analysis Toolkit (GATK) (DePristo et al., 2011). The resulting BAM files were analyzed to identify variants using the HaplotypeCaller module of GATK for multi-sample variant callings (Nho et al., 2014).

### Homology modeling

The model of the PLCG2 structure containing all residues from amino acid 14–1190 was built using the template PLCG1 model (Hajicek et al., 2019). The cartoons with substitutions of M28L and P522R were generated with PyMol (The PyMOL Molecular Graphics System, Version 2.0 Schrödinger, LLC).

### Bulk RNA-Seq analysis in the postmortem human brain

RNA-Seq data (logCPM normalized) were downloaded from the AMP-AD Consortium, in which individuals were participants in the Religious Orders Study and Memory and Aging Project (ROS/MAP) cohort (Bennett et al., 2018; Tsai et al., 2021). The procedures for sample collection, postmortem sample descriptions, tissue and RNA preparation methods, library preparation and sequencing methods, and sample quality controls were previously described in detail (De Jager et al., 2018).

### Differential gene expression and pathway enrichment analysis

The *limma* package in R software (Ritchie et al., 2015) was used to identify differentially expressed genes and to perform differential expression analyses of bulk RNA-Seq data from different samples. The ClusterProfiler package was used to automate biological-term classification and enrichment analysis for the differentially expressed genes (Yu et al., 2012).

### Mice

The 5xFAD amyloid AD mouse model was used for immunofluorescence, cytokine production, qPCR and differential expression analyses. Mice were maintained on the C57BL/6J (B6) background and purchased from the Jackson Laboratory (JAX MMRRC Stock# 034848). The 5xFAD transgenic mice overexpress the following five FAD mutations under control of the Thy1 promoter: the APP (695) transgene containing the Swedish (K670N, M671L), Florida (I716V), and London (V7171) mutations, and the PSEN1 transgene containing the M146L and L286V FAD mutations (Oakley et al., 2006).

We also used two PLCG2^P522R^ and PLCG2^M28L^ mouse models recently generated by the IU/JAX/UCI MODEL-AD consortium (JAX MMRRC Stock# 029598 and #030674; https://www.model-ad.org/strain-table/). PLCG2^P522R^ and PLCG2^M28L^ mice were generated using CRISPR/cas9 endonuclease-mediated genome editing to introduce the mutations. The *APOE4* gene sequence and TREM2^R47H^ mutation from the PLCG2^M28L^ mouse model (Oblak et al., 2022) were moved by crossing with B6 mice. These mice were maintained on the B6 background and crossed with 5xFAD mice to yield the 5xFAD; PLCG2^M28L^ and 5xFAD; PLCG2^P522R^ genotypes (5xFAD^M28L^ and 5xFAD^P522R^). The same numbers of male and female mice were used in the current study.

Up to five mice were housed per cage with SaniChip bedding and LabDiet® 5K52/5K67 (6% fat) feed. The colony room was kept on a 12:12 h light/dark schedule with the lights on from 7:00 am to 7:00 pm daily. The mice were bred and housed in specific-pathogen-free conditions. Both male and female mice were used, and the numbers of male and female mice were equally distributed. The number of mice used for each experiment is stated in the corresponding figure legends, and the results of individual values are shown in the scatter plot.

Mice were euthanized by perfusion with ice-cold phosphate-buffered saline (PBS) following full anesthetization with Avertin® (125-250 mg/kg intraperitoneal injection). Animals used in the study were housed in the Stark Neurosciences Research Institute Laboratory Animal Resource Center at Indiana University School of Medicine. All animals were maintained, and experiments were performed, in accordance with the recommendations in the Guide for the Care and Use of Laboratory Animals of the National Institutes of Health. The protocol was approved by the Institutional Animal Care and Use Committee (IACUC) at Indiana University School of Medicine.

### Immunoblotting

Tissue was extracted and processed as described above, then centrifuged. Protein concentrations were measured with a BCA kit (Thermo Scientific). One hundred micrograms of protein per sample was denatured by heating the samples for 10 min at 95°C. The samples were then loaded into 4–12% Bis-Tris gels (Life Technologies) and run at 100 V for 90 min. The following primer antibodies were used: PLCG2 (CST #3872 1:500, Rabbit mAb) and β-Actin (Santa Cruz #sc-47778). Each sample was normalized to β-Actin, and the graphs represent the values normalized to the mean of the WT mouse group at each time point.

### Immunofluorescence and image analysis

Perfused brains from mice at 7.5 months of age were fixed in 4% paraformaldehyde for 24 h at 4°C. Following 24 h fixation, brains were cryoprotected in 30% sucrose at 4°C and embedded. Brains were cut on a microtome into 30-μm free-floating sections. For immunostaining, at least three matched brain sections were used. Free-floating sections were washed and permeabilized in 0.1% TritonX in PBS (PBST), and antigen retrieval was subsequently performed using 1x Reveal Decloaker (Biocare Medical) at 85°C for 10 min. Sections were blocked in 5% normal donkey serum in PBST for 1 h at room temperature (RT). The sections were then incubated with the following primary antibodies in 5% normal donkey serum in PBST overnight at 4°C: Iba1 (Novus Biologicals #NB100-1028, goat, 1:1000); 6E10 (BioLegend #803001, mouse, 1:1000; AB_2564653); NeuN (Abcam, ab104225, rabbit, 1:1000); CLEC7A (InvivoGen, mabg-mdect, rat, 1:500 of 1 mg/ml); and P2RY12 (AnaSpec, AS-55043A, rabbit, 1:1000). Sections were washed and visualized using the respective species-specific AlexaFluor fluorescent antibodies (diluted 1:1000 in 5% normal donkey serum in PBST for 1 h at RT). Sections were counterstained with antibodies and mounted onto slides. For X34 staining (Sigma, #SML1954, 100 uM), sections were dried at RT, rehydrated in PBST, and stained for ten mins at RT. Sections were then washed five times in double-distilled water and washed again in PBST for five mins (Styren et al., 2000). Images were acquired on a fluorescence microscope with similar exposure and gains across stains and animals.

Images were taken on a Leica DM6 microscope or a Nikon A1R confocal microscope (Nikon Instruments, Melville, NY) for higher magnification and analyzed using ImageJ software (NIH) (Schneider et al., 2012). The results were obtained from an average of at least three sections per mouse, with a threshold applied across all images. The threshold function in ImageJ was used to determine the percentage of the immunoreactive area (X34, 6E10, IBA1, PR2Y12 and CLEC7A) within the region of interest (ROI). Plaque categorization was performed by the quantification of the 6E10, X34 and 6E10+/X34+ costained area. For quantification of the staining percentage area of X34 plaques, the surface of X34 plaques was created by the create selection function, and the percentage of the immunoreactive area was analyzed within the X34 positive extended surface. Plots were generated using GraphPad Prism (Version 9.3.1).

### Magnetic resonance imaging

High-resolution T2*-weighted magnetic resonance (MR) images were acquired using a Bruker BioSpec 9.4T/30 MRI scanner outfitted with a high-sensitivity cryogenic RF surface receive-only coil (Bruker CyroProbe) as previously described (Maharjan et al., 2022). Images were acquired using a 3D gradient echo sequence with the following acquisition parameters: TR: 200 ms; TE: 10 ms; Ave: 2; Flip Angle: 45; Spatial resolution 25 μm isotropic. To assess amyloid deposition in the brain of AD mice, ITK-SNAP (Yushkevich et al., 2006) was used to determine the hypo-intense area in MR images.

### Primary microglia culture

Microglia were isolated as previously described (Saura et al., 2003). Briefly, brain tissue from C57BL/6 neonatal mice aged P2–P4 was homogenized in Dulbecco’s Modified Eagle Medium (DMEM) (Gibco™ 10566016), filtered through 250 and 100 μM meshes sequentially, and cultured in Advanced DMEM/F12 (Gibco™ 12634028) supplemented with 10% FBS (Gibco™ 16000-044), 2 mM L-glutamine (Gibco™ 25030081) and Penicillin/Streptomycin (Gibco™ 15140122). After 21 days in culture, the cells were subjected to mild trypsinization using a 1:3 dilution of EDTA-Trypsin (Gibco™ 25-200-072) in DMEM for 20 min. Trypsinization resulted in the detachment of an intact layer of astrocytes, leaving microglia attached to the bottom of the plate, which were used for experiments within 48 h. For the RNA analysis, microglia were cultured in polystyrene 6-well plates (Falcon™ 08-772-1B); for immunofluorescence, microglia were cultured in a chamber slide system (Nunc™ Lab-Tek™ II Chamber Slide™, 154453).

### Aβ_1-42_ aggregation and uptake assay

The HiLyte™ Fluor 555-labeled Aβ_1-42_ peptide was obtained from AnaSpec (AS-60480-01), reconstituted as suggested by the manufacturer with 1.0% ammonium hydroxide and phosphate-buffered saline (PBS) pH=7.4 up to 1 mg/ml. The peptides were diluted to 0.5 mg/ml in PBS pH=7.4 and aggregated at 37°C for 5 days, similar to a previously described method (Moutinho et al., 2022). Incubation of microglia with HiLyte™ Fluor 488-labeled Aβ_1-42_ aggregates was followed by the immunofluorescence protocol.

### Y-maze task

Working memory in mice was assessed by spontaneous alternation in the Y-maze task as previously described (Moutinho et al., 2022). Animals were placed in the Y-maze with free access to all three arms and tracked and recorded for 8 min using ANY-maze software (Stoelting Co.). The number of alternations was determined by counting the sequential entries into the three different arms of the maze (same-arm re-entries were possible). The percentage of spontaneous alternation was calculated using the following formula: (number of alternations performed)/(number of possible alternations [total arm entries-2])x100. An entry counted when all four limbs of a mouse entered the arm.

### Hippocampal Slice Electrophysiology

Mice were deeply anesthetized and intracardially perfused with ice-cold artificial cerebrospinal fluid (aCSF) containing (in mM) 124 NaCl, 4.5 KCl, 1.2 NaH_2_PO_4_, 26 NaHCO_3_, 10 glucose, 1 MgCl_2,_ and 2 CaCl_2,_ continuously bubbled with 95% O_2_/5% CO_2_; pH 7.4, 310 mOsm. The brains were quickly removed, and hippocampal slices (280 µm) were cut at 0.1 mm/s with a Leica VT1200 vibratome in an ice-cold oxygenated sucrose-based solution containing (in mM) 194 sucrose, 10 glucose, 30 NaCl, 26 NaHCO_3_, 0.5 NaH_2_PO_4_,4.5 KCl, and 1 MgCl_2,_ pH 7.4. After 60 min recovery in an incubation chamber containing oxygenated aCSF solution at 33°C, the slices were then kept at room temperature until they were transferred to a recording chamber, perfused continuously (∼2 ml/min) with oxygenated aCSF at ∼32°C.

### Field recordings

Field excitatory postsynaptic potentials (fEPSP) were recorded with micropipettes filled with 1 M NaCl placed in the stratum radiatum of the CA1 region of the hippocampus. A stimulating stainless steel stereotrode (1 MΩ) was placed in the Schaffer collateral pathway. The intensity of the stimulator was increased stepwise until a maximal response was obtained using a constant current isolated stimulator (Digitimer). Signals were acquired using a Multiclamp 700B amplifier and Clampex software (Molecular Devices). Signals were low pass filtered and digitized at 50 kHz, and the slope of the fEPSP (mV/ms) was measured. Paired-pulse ratios (PPR) were obtained every 20 s at 40 ms increasing inter-stimuli interval (ISI). For LTP recordings, the following protocol was used: after adjusting the stimulation strength to produce 50% of the maximum intensity, a stable 10 min of baseline (30 pulses every 20 s) was recorded, followed by 1 min conditioning trains (10 pulses at 100Hz) repeated 4 times every 20 s; a 60 min post-conditioning was performed at the same baseline stimulation frequency. The synaptic strength change was expressed as the percentage of change with respect to the average baseline.

### Whole-cell recordings

Patch pipettes (2.5–3.5 MΩ) were pulled (Sutter Instruments) from borosilicate glass (World Precision Instruments) and filled with an internal solution containing (in mM) 120 CsMeSO_3_; 5 NaCl, 10 TEA, 10 HEPES, 5 lidocaine bromide, 1.1 EGTA, 4 Mg-ATP, and 0.3 Na-GTP (pH adjusted to 7.2 and 290 mOsm). Neurons in the CA1 region were visualized with a 40X water-immersion objective with infrared-differential interference contrast video microscopy (BX51WI; Olympus). Spontaneous excitatory postsynaptic currents (sEPSC) were gap-free recorded five mins after breaking for 3 min in voltage-clamp mode with the membrane potential (V_h_) held at - 70 mV. sIPSCs were recorded with the membrane potential held at +10 mV. Recordings were conducted using a Bessel filter set at 4 kHz and digitized at 50 kHz with a Digidata 1440A A/D interface (Molecular Devices). A negative 5 mV pulse was delivered regularly to monitor access and input resistance. Recordings with access resistance greater than 25 MΩ or with changes in access or greater than 20% were discarded. Recordings of AMPA/NMDA ratios were performed after adding picrotoxin (50 µM) to the aCSF to block inhibitory currents. Evoked synaptic responses were evoked at V_h_=+40 mV before and after the addition of d-APV (50 μM) to block NMDA receptors and isolate AMPA currents. pClamp11 (Molecular Devices) and MiniAnalysis (Synaptosoft) were used for quantification. Statistical analyses were conducted using Prism (GraphPad Software). All data are presented as the mean ± SEM. The significance level was set at p<0.05

### Mouse RNA isolation for qPCR and NanoString nCounter analysis

Mice were anesthetized with Avertin and perfused with ice-cold PBS. The cortical and hippocampal regions were microdissected and stored at -80°C. Frozen brain tissues were homogenized in T-PER tissue protein extraction reagent (Catalog #78510, Thermo Fisher) and stored in an equal volume of RNA STAT-60 (Amsbio) at -80°C until RNA extraction was performed. RNA was isolated by chloroform extraction and purified using the Purelink RNA Mini Kit (Life Technologies). cDNA was prepared from 750 ng of RNA using the High-Capacity of RNA-to-cDNA kit (Applied Biosystems), and qPCR was performed on the StepOne Plus Real-Time PCR system (Life Technologies) with the Taqman Gene Expression Assay (Plcg2, Mm00549424_m1, Life Technologies). Relative gene expression was determined with the △△CT method and was assessed relative to Gapdh (Mm99999915_g1). Statistical analyses of the qPCR results were performed using a one-way analysis of variance (ANOVA) test followed by Tukey’s post hoc to compare genotypes (GraphPad Prism, version 9.3.1).

Glial Profiling and Neuropathology Panels were used for the NanoString nCounter analysis (NanoString Technologies, Seattle, WA, USA). Two hundred nanograms of RNA was loaded per 7.5-month-old male and female mouse from the 5xFAD, 5xFAD^M28L^ and 5xFAD^P522R^ mouse samples (*N*=12 per genotype, 6 male and 6 female mice) and hybridized with probes for 16 h at 65°C. The results obtained from the nCounter MAX Analysis System (NanoString Technologies, catalog #NCT-SYST-LS, Seattle WA) were imported into the nSolver Analysis Software (v4.0; NanoString Technologies) for QC verification, normalization, and data statistical analysis using Advanced Analysis software (v2.0.115; NanoString Technologies). All assays were performed according to the manufacturer’s protocols (Preuss et al., 2020; Reilly et al., 2020).

### Quantification of proinflammatory cytokine levels in mouse brains

Protein levels of inflammatory cytokines in the mouse brains were quantified using Meso Scale Discovery (MSD) 96-well multispot V-PLEX Proinflammatory Panel I (K15048D, MSD, Gaithersburg, MD, USA), following the manufacturer’s guidelines (Oblak et al., 2021). Frozen brain cortex samples were homogenized in T-PER tissue protein extraction reagent supplemented with protease and phosphatase inhibitor cocktails (Sigma-Aldrich). Total protein concentration was measured using the Pierce BCA Protein Assay Kit (Thermo Scientific). Fifty microliters of protein lysate (500 µg) was used to analyze the proinflammatory cytokine content (*N*=10 per genotype, 7.5-month-old, 5 male and 5 female mice).

### Nuclei isolation and fluorescence-activated cell sorting (FACS)

Nuclei were isolated as previously described (Habib et al., 2017; Hahn et al., 2021). Nuclei were isolated from the fresh-frozen cortical brain region of the left hemisphere. All reagents were placed on ice. Frozen tissue was minced with a chilled razor blade and then Dounce homogenized 25 times with a loose pastle A followed by 15 times with a tight pastle B (Sigma-Aldrich, St. Louis, USA #D8938), all while simultaneously twisting up and down with lysis buffer from the Nuclei EZ Prep Kit (Sigma-Aldrich, St. Louis, USA, #Nuc101-1KT). The tube was incubated on ice for 5 min. Nuclei were pelleted with 5 min centrifugation at 500 g (4°C). The pellet was then resuspended with 4 ml fresh lysis buffer. Following a subsequent 5 min centrifugation step at 500 g (4°C), the lysis buffer was removed. The pellet was washed with 4 ml PBS and then transferred into a FACS tube.

Nuclei were centrifuged for 10 min at 300 g (4°C) and resuspended in 50 μl FACS buffer (1% BSA, 1× PBS; sterile filtered) containing 2 U/ml Protector RNase Inhibitor (Sigma-Aldrich, St. Louis, USA, #3335402001) with anti-CD16/CD32 Fc Block (BD Biosciences #553142). After 5 min incubation, an additional 50 μl of antibody cocktail with Protector RNase Inhibitor was added: 1 μl CD45 (BD Biosciences #552848), 1 μl Olig2 (Millipore #MABN50A4), and 1 μl NeuN (Abcam #ab190565) in 50 μl FACS buffer. Following Fc blocking, 50 μl of antibody master mix was added to each sample to achieve 1× antibody concentration. Samples were incubated with the staining antibodies on ice with gentle shaking for 30 min and then centrifuged for 10 min at 300 g (4°C) before being resuspended in 700 μl of FACS buffer with Protector RNase Inhibitor. Subsequently, 1 µl Hoechst 33342 was added.

NeuN-positive and NeuN-negative nuclei were separately sorted into 1.5 ml DNA LoBind tubes with 1 ml Dijon buffer (200 μl UltraPure BSA, Thermo, #AM2618, 800 μl PBS, and 5 μl Protector RNase Inhibitor. A total of 50,000 single nuclei were sorted from each sample. Nuclei were counted, and concentrations were adjusted to ∼1 mio nuclei/ml.

### Chromium 10X library generation and Illumina sequencing

Reagents for the Chromium Single Cell 3ʹ Library & Gel Bead Kit v3 (10X Genomics, Pleasanton, USA) were thawed and prepared according to the manufacturer’s protocol. The nuclei/master mix solution was adjusted to target 10,000 nuclei per sample (5,000 nuclei from male mice and 5,000 nuclei from female mice) and loaded on a standard Chromium Controller (10X Genomics, Pleasanton, USA), according to the manufacturer’s protocol. All reaction and quality control steps, including library construction (using Chromium Single Cell 3ʹ Library Construction Kit v3), were conducted according to the manufacturer’s protocol and with the recommended reagents, consumables and instruments. Quality control of cDNA and libraries was conducted using a Bioanalyzer (Agilent, Santa Clara, USA) at the Stanford Protein and Nucleic Acid Facility.

Illumina sequencing of 10X snRNA-seq libraries was performed by Novogene Co. Inc. (Sacramento, USA; https://en.novogene.com/). Multiplexed libraries were sequenced with 2 × 150-bp paired-end (PE) reads in a single S4 lane on an Illumina Novaseq S4 (Illumina, San Diego, USA) targeting 100 million reads per library. Novogene conducted base-calling, demultiplexing, and the generation of FastQ files.

## Resource availability

### Lead contact

Further information and requests for resources and reagents should be directed to and will be fulfilled by lead contact Gary E. Landreth.

### Materials availability

This study did not generate new unique reagents.

### Data and code availability

All data supporting the findings of the article are available within the main text or supplemental information. All original code will be shared by the lead contact upon request. Any additional information required to reanalyze the data reported in this paper is available from the lead contact upon request.

## Acknowledgments

The authors thank the members of the Landreth and Lamb laboratory for feedback and support throughout the study. We thank Louise Pay for her critical comments on the manuscript and Teaya N. Thomas for the help with taking care of the mice. This work was supported by NIA grant K01 AG054753 (A.L.O), NINDS grant R01 NS125020 (N.W), NIA grant R03 AG063250 (K.N), NIH grant NLM R01 LM012535 (K.N), NIA grant U54 AG054345 (B.T.L et al.), and NIA grant RF1 AG074566 (B.T.L, S.J.B, and G.E.L).

## Authors’ contributions

A.P.T, C.D, P.B.L, G.V.D.P, A.J.C, E.K.L, N.H, O.H, M.A, A.G.F, N.W, J.P, G.X, and K.N conducted experiments, collected, and analyzed data. A.P.T, A.L.O, K.N, B.T.L. and G.E.L conceptualized and designed experiments. A.P.T, C.D, P.B.L, G.V.D.P, A.C, E.K.L, N.H, M.M, T.W.C, B.K.A, J.S, Q.Z, A.D.M, Y.L, S.J.B, K.N, B.T.L, and G.E.L discussed and interpreted data. A.P.T and G.E.L designed figures and wrote the manuscript. G.E.L supervised, directed, and managed the study. All authors discussed the results and commented on the manuscript.

## Inclusion and diversity

We support inclusive, diverse, and equitable conduct of research. One or more of the authors of this paper self-identifies as an underrepresented ethnic minority in science. One or more of the authors of this paper self-identifies as a gender minority in their field of research.

## Supplemental information

**Figure S1.**
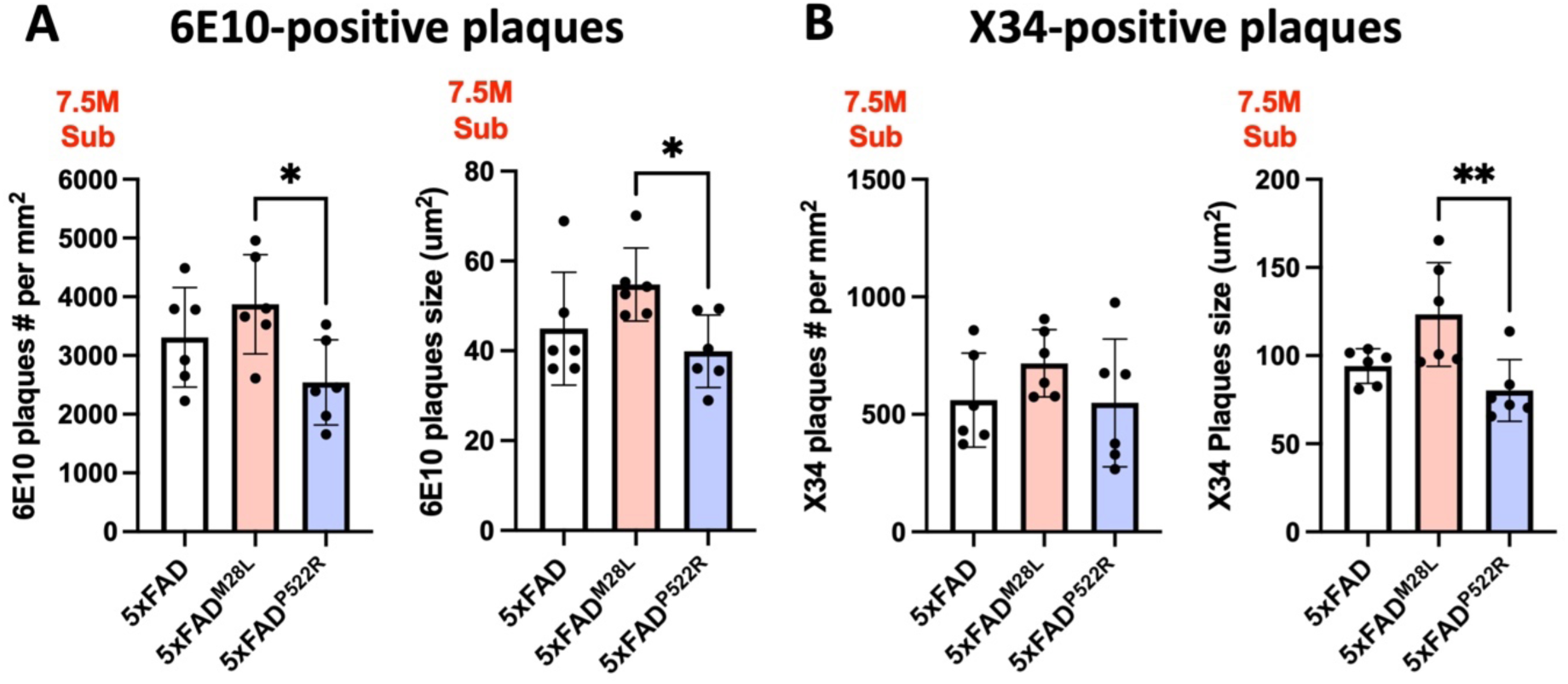
Numbers and sizes of 6E10-positive and X34-positive plaques in AD mice **(A)** Scatter plots show the quantification of the numbers and sizes of 6E10-positive plaques. **(B)**Scatter plots show the quantification of the numbers and sizes of X34-positive plaques.

**Figure S2.**
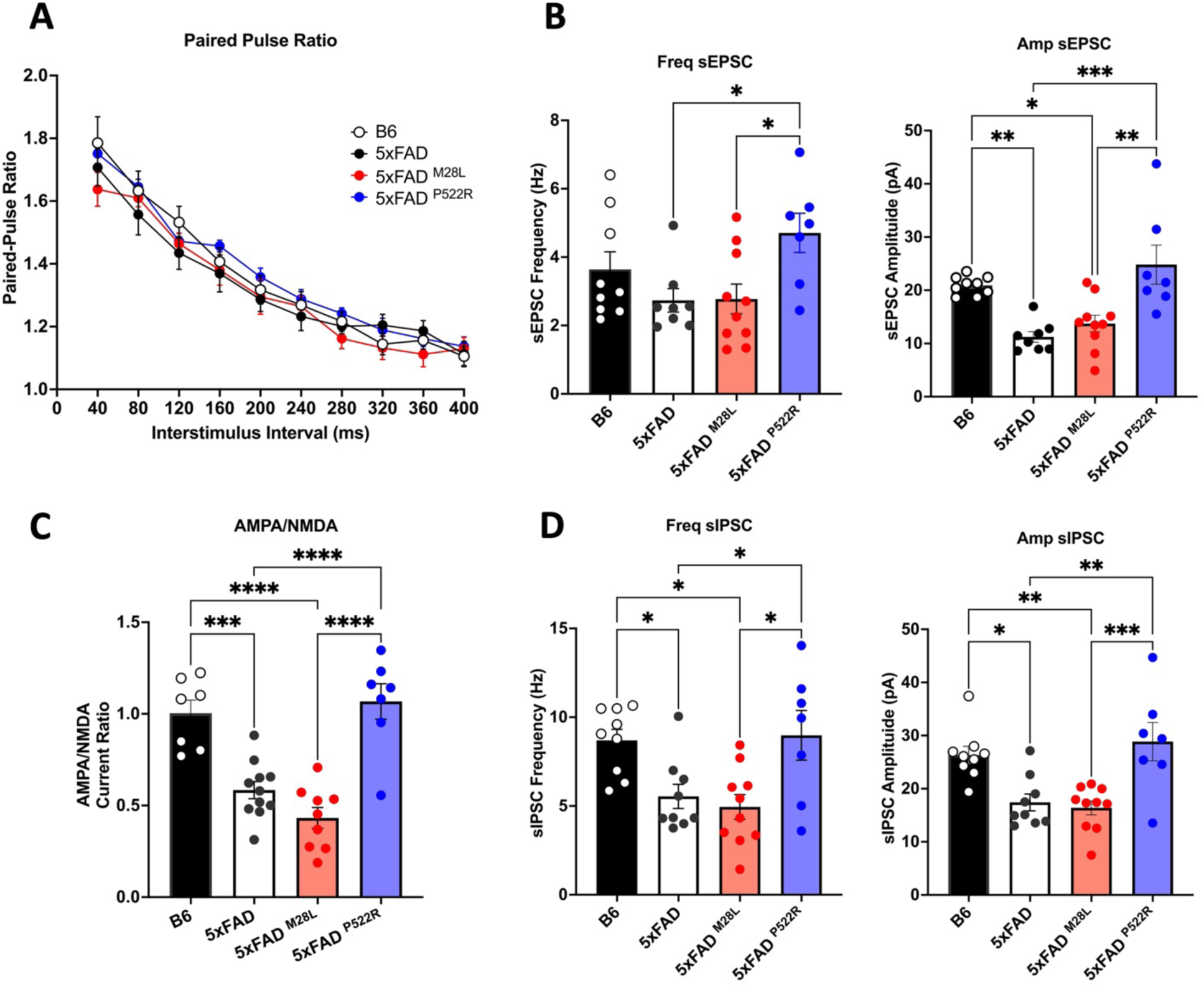
Paired-pulse ratios and whole-cell recordings in AD mice **(A)** Paired-pulse ratios were obtained every 20 s at 40 ms increasing inter-stimuli intervals, and no genotype differences were observed. **(B-D)** Whole-cell patch-clamp electrophysiological recordings were performed on hippocampal area CA1 pyramidal neurons. **(B)** Measures of spontaneous excitatory postsynaptic currents (sEPSCs) parameters (frequency and amplitude). **(C)** Measures of differences in excitatory transmission by measuring EPSCs mediated by AMPA and NMDA glutamate receptors. **(D)** Measures of sIPSC parameters (frequency and amplitude). The software pClamp11 (Molecular Devices) and MiniAnalysis (Synaptosoft) were used for quantification. Statistical analyses were performed using Prism (GraphPad Software). All data are presented as the mean ± SEM. * P <0.05; ** P <0.01; *** P <0.001; **** P <0.0001.

